# Consistent signatures of urban adaptation in a native, urban invader ant *Tapinoma sessile*

**DOI:** 10.1101/2021.06.21.449338

**Authors:** Alexander J. Blumenfeld, Pierre-André Eyer, Anjel M. Helms, Grzegorz Buczkowski, Edward L. Vargo

**Affiliations:** Department of Entomology, 2143 TAMU, Texas A&M University, College Station, Texas, 77843-2143, USA; Department of Entomology, Purdue University, West Lafayette, Indiana, 47907, USA

**Author notes:** Correspondence should be addressed to A.J.B.

## Abstract

Biological invasions are becoming more prevalent due to the rise of global trade and expansion of urban areas. Ants are among the most prolific invaders, with many exhibiting a multi-queen colony structure, dispersal through budding and a lack of inter-nest aggression. Although these characteristics are generally associated with the invasions of exotic ants, they may also facilitate the spread of native ants into novel habitats (*e*.*g*., urban areas). Native to North American forests, the odorous house ant *Tapinoma sessile* has become abundant in urban environments throughout the United States. Forest-dwelling colonies typically have a small workforce, inhabit a single nest, and are headed by a single queen, whereas urban colonies tend to be several orders of magnitude larger, inhabit multiple nests and are headed by multiple queens. Here, we explore and compare the population genetic and breeding structure of *T. sessile* within and between urban and natural environments in several localities across its distribution range. We found the social structure of a colony to be a plastic trait in both habitats, although extreme polygyny (*i*.*e*., nests with multiple queens) was confined to urban habitats. Additionally, polydomous colonies (*i*.*e*., nests lacking genetic differentiation and behavioral antagonism) were only present in urban habitats, suggesting *T. sessile* can only achieve unicoloniality within urbanized areas. Finally, we identified strong differentiation between urban and natural populations in each locality and continent-wide, indicating cities may restrict gene flow and exert intense selection pressure. Overall, our study highlights urbanization’s influence in charting the evolutionary course for species.

## Introduction

Biological invasions are becoming more prevalent throughout the world due to the rise of global trade (Banks, Paini, Bayliss, & Hodda, 2015) and the expansion of urban areas (Seto, Fragkias, Guneralp, & Reilly, 2011). This increasingly inter-connected world has allowed many species of plants and animals to become established and invasive within novel habitats (Meyerson & Mooney, 2007). Species invasions may promote ecological disturbance, whereby invasive species outcompete native species for resources or fill an empty ecological niche and alter the complexion of an ecosystem (Ehrenfeld, 2010). Ants are among the most prolific of these invaders (Jeschke & Wittenborn, 2011), with around 240 invasive species globally (Bertelsmeier, Ollier, Liebhold, & Keller, 2017). It is generally assumed that individual and colony-level plasticity in physiology, morphology and behavior may enhance their invasion success. This plasticity may allow for a rapid acclimation to novel ecological pressures faced in the introduced range to quickly impose their ecological dominance.

Many invasive ants possess a suite of shared characteristics that facilitate their success within new environments, such as a polygyne social structure (*i*.*e*., multiple reproductive queens), dispersal through budding and lack of aggression among non-nestmate workers (Eyer & Vargo, 2021; Lester & Gruber, 2016; McGlynn, 1999; Tsutsui & Suarez, 2003). A polygyne social structure may enhance the survival rate of a colony, as the colony can withstand the death of a single queen or multiple queens. The reproduction of multiple queens also allows for greater colony growth due to an increased production of new workers (Boomsma, Huszár, & Pedersen, 2014; Boulay, Arnan, Cerdá, & Retana, 2014). Oftentimes, polygyny is associated with colony foundation through budding (Cronin, Molet, Doums, Monnin, & Peeters, 2013). This dispersal strategy entails the founding of new colonies by queens assisted by workers, dispersing from their natal nests on foot to establish new nests nearby (Hölldobler & Wilson, 1977; Keller, 1991). This strategy is associated with high foundation success, as the help of workers increase survival and reproduction of new nests during the early establishment stage (Cronin et al., 2013). However, this mode of foundation restricts dispersal of the species, often leading to the establishment of many genetically similar colonies across a landscape and thereby a pattern of isolationby-distance (Schultner, Saramäki, & Helanterä, 2016). Polygyny and dispersion through budding may increase the chance of invasion and spread, as they may increase the propagule pressure of a species (*i*.*e*., the number of reproductive individuals introduced and frequency of introduction events) (Tsutsui & Suarez, 2003; Yang et al., 2012).

In most invasive ants, invasion success is also enhanced by the lack of aggression between workers from distinct nests within invasive ranges. This absence of aggression facilitates the formation of supercolonies – extensive colonies comprised of many nests (*i*.*e*., polydomy) exchanging workers, queens, brood and resources (Tsutsui & Suarez, 2003). This colony structure eliminates intraspecific competition, leading to dense networks of interconnected nests, genetically indistinguishable from each other. The development of highly polygynous supercolonies enables invasive populations to reach tremendous densities and rapidly outcompete native species by allocating a high number of workers to monopolize resources (Tsutsui & Suarez, 2003). For example, populations of the yellow crazy ant *Anoplolepis gracilipes* and the little fire ant *Wasmannia auropunctata* can reach densities of up to 20 million and 240 million ants per hectare, respectively (Abbott, 2005; Souza, Follett, Price, & Stacy, 2008). To date, the association of polygyny, dispersion via budding and development of a dense polydomous nest structure have been observed in many invasive ants, such as *Linepithema humile, Pheidole megacephala, Monomorium pharaonis* and *Nylanderia fulva* (Buczkowski & Bennett, 2009; Eyer et al., 2018; Tsutsui, Suarez, Holway, & Case, 2000).

Three major hypotheses have been suggested to explain the formation of supercolonies in introduced populations (Giraud, Pedersen, & Keller, 2002; Steiner et al., 2007). First, because nestmate recognition is partly based on heritable cues, decreased genetic diversity in introduced populations as a result of a genetic bottleneck reduces recognition templates among colonies (Beye, Neumann, Chapuisat, Pamilo, & Moritz, 1998; Lockey, 1991; Pirk, Neumann, Moritz, & Pamilo, 2001). Ultimately, the loss of diversity at the recognition locus (or loci) may homogenize recognition templates among colonies, thereby preventing workers from discriminating against non-nestmate conspecifics (Tsutsui, Suarez, & Grosberg, 2003; Tsutsui et al., 2000; Vásquez, Schal, & Silverman, 2008). A second hypothesis proposes that reduced nestmate recognition was selected for to avoid repeated fights with conspecifics in the densely populated introduced range (Chapuisat, Bernasconi, Hoehn, & Reuter, 2005; Giraud et al., 2002), whereby nonaggressive colonies reach larger sizes and outcompete aggressive ones (Holway, 1999; Holway, Suarez, & Case, 1998; Katzerke, Neumann, Pirk, Bliss, & Moritz, 2006). The last hypothesis suggests that polydomy and polygyny are pre-existing characteristics of the species in its native range (Giraud et al., 2002; Pedersen, Krieger, Vogel, Giraud, & Keller, 2006). The lack of conspecific competition in the introduced range may have allowed colonies of species that are polygynous and polydomous to grow extremely large.

The odorous house ant *Tapinoma sessile* is a native, urban-invader that exhibits a similar transition in its breeding system and social structure between its natural and urban populations. Native to the temperate deciduous forests of North America, this ant has become highly abundant in urban environments throughout the United States (Buczkowski, 2010; Buczkowski & Bennett, 2008; Menke et al., 2010). Though forest-dwelling colonies are still prevalent, the life history traits of colonies in the two habitats are vastly different. Forest-dwelling colonies are typically small (<200 workers) and consist of a single nest headed by a single queen (Buczkowski, 2010). On the other hand, urban colonies tend to be large (>100,000 workers) and made of several interconnected nests, each comprising numerous reproductive queens, with low inter-nest aggression over large landscapes. This suggests the existence of supercolonies in this species within urban environments (Buczkowski, 2010; Buczkowski & Bennett, 2008; Menke et al., 2010). However, these assessments have only been based on behavioral studies (Buczkowski, 2010; Buczkowski & Bennett, 2008; Buczkowski & Krushelnycky, 2011), while the genetic underpinnings of the colonies have not been analyzed. In *T. sessile*, four major mitochondrial clades have been described across the United States (Menke et al., 2010). Remarkably, this shift in life history traits has occurred consistently across its distribution, rather than all urban colonies originating from a single forest population (Menke et al., 2010). Therefore, plasticity in colony structure appears to be inherent within the species. This repeated transition of small, monogyne forest-dwelling colonies to large, polygyne urban colonies resembles the invasions of exotic ants into novel environments. Thus, *T. sessile* represents a unique opportunity to determine the factors driving these trait differences, which may provide insights into their evolutionary trajectory and broaden our understanding of the mechanisms linking them to species invasions.

Here, we conducted a large-scale analysis of the population and colony structure of *T. sessile* across the four geographic clades uncovered within the United States (Menke et al., 2010). For each of the four clades, we performed a paired sampling of one urban and one natural habitat in close geographic proximity (except for the Mountain clade in Colorado – see Methods). We first investigated the breeding structure of these populations to test for consistent transitions of monogyne colonies in natural habitats to polygyne colonies in urban areas, by assessing the number of queens per nest and the relatedness among nestmate workers. We then evaluated the colony structure of *T. sessile* in each locality by genetically inferring whether different nests belong to the same polydomous colony, testing for unicoloniality within urban habitats and multicoloniality within natural habitats. We also analyzed whether workers from different nests recognize each other as colony-mates through behavioral assays, testing for reduced aggression between non-nestmate workers in urban habitats compared to natural habitats, and assessed whether this discrimination toward non-nestmate workers is mediated through chemical cues. In addition, we investigated the dispersal ability of *T. sessile* by testing for an isolation-by-distance pattern in each locality and habitat. Finally, we discuss the potential evolutionary mechanisms enabling urban invasions by a native ant species, comparing these mechanisms with the life history traits shared by most invasive ant species.

## Materials and Methods

### Study sites and sampling

Nests of *T. sessile* were collected from July of 2018 to August of 2020 in four localities across the United States: Bloomington, Indiana; Bay Area, California; Little Rock, Arkansas; and Boulder, Colorado (Figure 1a). These four localities correspond to the four geographic clades previously elucidated by Menke et al. (2010). For each locality, two sites in close geographic proximity were identified – one comprising the urban environment (residential or commercial areas) and the other comprising the natural environment (forests largely free of anthropogenic influence), with fifteen nests collected in each habitat. No nests were found in the urban environment of Boulder, CO. Therefore, our total collection consisted of 104 nests across the four localities and seven total sites (details of the sampling are given in Table S1). Entire nests were sampled to ensure a reliable count of queens and that ants collected belonged to the same nest. The nests were transported to the laboratory and kept under standard conditions (28 ± 2°C, 12:12 h light period, and fed with an artificial ant diet (Dussutour & Simpson, 2008)). For each nest, eight workers were separately placed in 200µL hexane for chemical analysis, while a subset of workers, queens and males were directly stored in 96% ethanol at 4°C for genetic analysis.

**Figure 1:**
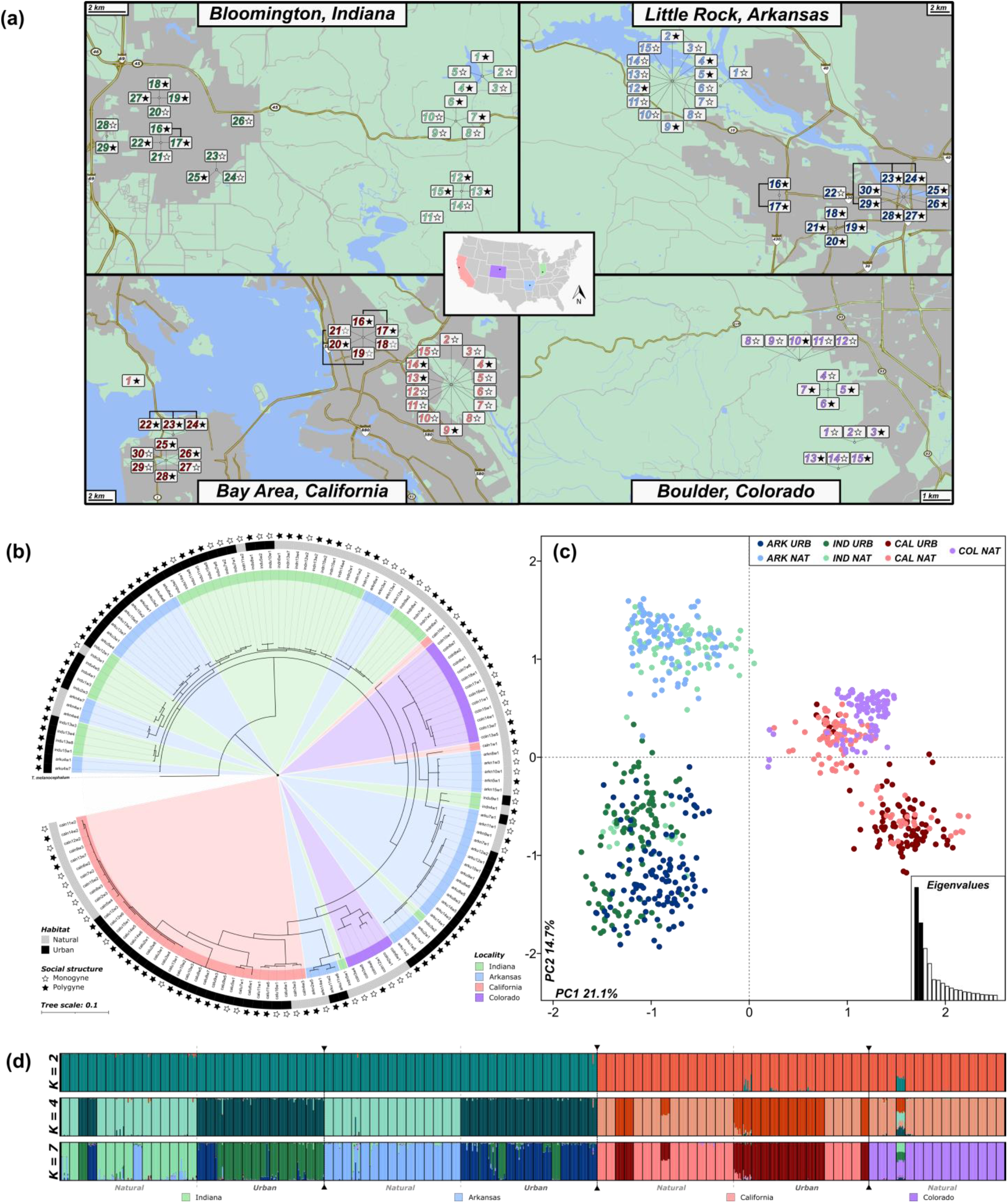
**(a)** Sampling locations of *Tapinoma sessile* across the United States. For each locality, nests were sampled in both natural and urban environments, depicted as light-colored (natural) and dark-colored (urban) numbers in the figure. The thickened black lines connecting some nests represent nests that were found to belong to the same colony. Additionally, the stars next to each number denote the social structure of the colony - white and black stars represent monogyne and polygyne colonies, respectively. Note that for each locality, the counting always begins with the first natural nest; also, note that no urban nests were found in Colorado. **(b)** Bayesian inference tree based on 145 COI sequences of *T. sessile* across the four localities, with one *T. melanocephalum* sequence as an outgroup. **(c)** PCA of each individual from each nest sampled in the overall dataset (dots represent individuals). **(d)** STRUCTURE analysis across four values of *K*, which correspond to the levels of hierarchy present within the overall dataset (2 = habitat; 4 = locality; 7 = habitat x locality).

### Genetic Analyses

The genomic DNA of eight workers and up to eight queens and males from each nest was extracted following a modified Gentra Puregene extraction method (Gentra Systems, Inc. Minneapolis, MN, USA). Species-specific microsatellite primers do not exist for *T. sessile*; instead, we tested 39 markers shown to amplify in closely related species (Berman, Austin, & Miller, 2014; Butler, Siletti, Oxley, & Kronauer, 2014; Krieger & Keller, 1999; Zheng, Yang, Zeng, Vargo, & Xu, 2018; Zima, Lebrasseur, Borovanska, & Janda, 2016). Forward primers were affixed with an M13 tail to enable PCR multiplexing via fluorescent labelling with 6-FAM, VIC, PET, and NED (Boutin-Ganache, Raposo, Raymond, & Deschepper, 2001). PCR conditions and multiplexing arrangements are given in the online supplementary material (Table S2). PCR reactions were performed on a Bio-Rad thermocycler T100 (Bio-Rad, Pleasanton, CA, U.S.A.). Multiplex PCR products were run on an ABI 3500 capillary sequencer (Applied Biosystems, Foster City, CA, U.S.A.) along with the LIZ500 standard. Geneious v.9.1 was used for scoring alleles (Kearse et al., 2012). Of the 39 markers tested, 21 were discarded due to non-amplification or monomorphic amplification. The linkage disequilibrium (LD) for each pair of loci was tested for each locality separately using GENEPOP v4.7 (Rousset, 2008), with *p*-values corrected via the Holm method to account for multiple comparisons (Holm, 1979). Loci exhibiting linkage disequilibrium were discarded from further analysis. Overall, the final dataset includes 831 workers genotyped at 12 polymorphic microsatellite loci.

Sequencing of the cytochrome oxidase 1 (COI) mitochondrial gene was performed on at least one worker from each nest, with multiple workers from a nest sequenced if genotypes suggested they originated from different queens (n = 145). Gene sequences were amplified using primers LepF1 and LepR1, targeting a 658-bp fragment (Hajibabaei, Janzen, Burns, Hallwachs, & Hebert, 2006; Hebert, Penton, Burns, Janzen, & Hallwachs, 2004). PCR products were purified with EXOSAP-it PCR purification kit (Affymetrix) and sequenced using the ABI BigDye Terminator v.3.1 Cycle Sequencing Kit on an ABI 3500 Genetic Analyzer (Applied Biosystems). Base calling and sequence reconciliation were performed using CodonCode Aligner (CodonCode Corporation, Dedham, MA, USA).

### Population and colony structure analyses

#### Mitochondrial dataset

A Bayesian phylogenetic tree and a haplotype network were constructed to assign each nest into one of the four clades previously described by Menke et al. (2010). In addition, the mitochondrial dataset was used to test for the presence of multiple haplotypes within nests, which would indicate the reproduction of multiple unrelated queens. MrBayes v.3.2 was used to construct the tree (Ronquist et al., 2012), using the generalized time reversible model with gamma-distributed rate variation across sites and a proportion of invariable sites as the evolutionary model. Two simultaneous MCMC simulations ran for 2 × 10^6^ generations using four chains (three heated and one cold), with each run sampled every 500 generations. The mitochondrial network was produced via the median-joining method (Röhl, Bandelt, & Forster, 1999) implemented in POPART (Leigh & Bryant, 2015). The COI gene was extracted from the complete mitochondrial genome sequence of T. melanocephalum to use as an outgroup for both analyses (Du, Song, Yu, & Lu, 2019). Additionally, sequence divergence was compared within and between habitats and populations of T. sessile using the Kimura 2-parameter model (Kimura, 1980) in MEGA v. 10.2.2 (Kumar, Stecher, Li, Knyaz, & Tamura, 2018; Tamura et al., 2011).

#### Microsatellite dataset

The number and frequency of alleles, F-statistics (Weir & Cockerham, 1984), and observed and expected heterozygosity (Nei, 1987) were calculated for each microsatellite marker, and for each locality and habitat (as well as overall values) using FSTAT v2.9.4 (Goudet, 2003). For the overall dataset, the hierarchical partitioning of the genetic diversity between localities, between habitats within localities, between nests within habitats, between individuals within nests, and within individuals was assessed using an analysis of molecular variance (AMOVA) implemented in the ade4 R package (Dray & Dufour, 2007; R Core Team, 2020) via Poppr (Kamvar, Tabima, & Grünwald, 2014).

Three complementary approaches were used to determine whether workers collected from different nests belonged to the same colony. First, genotypic differentiation between each pair of nests within localities was tested using the log-likelihood G test implemented in GENEPOP v.4.7 (Rousset, 2008). Nests were considered distinct colonies if genotypic differentiation was found to be significantly different using Fisher’s test together with the Holm method to account for multiple comparisons (Holm, 1979). Second, population structure was visualized with a principal component analysis (PCA) using the adegenet R package (Jombart, 2008; R Core Team, 2020). Third, the presence of genetic structure was tested using the Bayesian clustering method implemented in STRUCTURE v.2.3.4 (Falush, Stephens, & Pritchard, 2003; Pritchard, Stephens, & Donnelly, 2000). Simulations were run for K (i.e., genetic clusters) ranging from 1 to the maximum number of nests per dataset, with each run of K replicated 20 times. The analyses were run under the admixture model with correlated allele frequencies enabled. Each run was initiated with a 50,000 burn-in period, followed by 100,000 iterations of the MCMC. The most likely number of genetic clusters (K) was inferred using the methods of both Evanno, Regnaut, and Goudet (2005) and Puechmaille (2016), with the output visualized via CLUMPAK (Kopelman, Mayzel, Jakobsson, Rosenberg, & Mayrose, 2015), as implemented in the web-based software StructureSelector (Li & Liu, 2018). The method of Puechmaille (2016) aims at unraveling finer partitioning in the data, whereas the Evanno et al. (2005) method aims at describing the primary partitioning. Finally, isolation-by-distance analyses were performed with Mantel tests using the vegan R package (Oksanen et al., 2020; R Core Team, 2020), between matrices of genetic differentiation (F_ST_) and geographic distance. The PCA, STRUTURE and isolation-by-distance analyses were first performed for the overall dataset, then for each locality, and finally for each habitat within each locality.

### Breeding structure analyses

We estimated the number of queens per nest and the genetic relatedness among nestmate workers for each locality and habitat. We also explored the possibility that queens use thelytokous parthenogenesis for the production of new queens, as this strategy was previously reported in several invasive ant species (Fournier et al., 2005; Pearcy, Goodisman, & Keller, 2011; Rabeling & Kronauer, 2013). The presence (or lack thereof) of multiple queens per nest was first determined directly from field observations. For each nest, polygyny was confirmed genetically through the presence of multiple mitochondrial haplotypes and through the composition of worker microsatellite genotypes. Polygyny was inferred when worker genotypes could not be reliably assigned to a single queen (all workers carrying one of the two alleles of the mother queen at all microsatellite markers).

Relatedness coefficients (r) among nests were estimated using COANCESTRY v.1.0.1.9 (Wang, 2011), following algorithms described by Queller & Goodnight (1989). As differences in allele frequencies may exist between localities, relatedness coefficients were calculated separately for each of the localities. Additionally, relatedness coefficients were estimated at three separate levels – 1) between workers, 2) between queens, and 3) between workers and queens. Finally, we evaluated whether queens produce new queens through thelytokous parthenogenesis by comparing the heterozygosity level and relatedness between castes in each locality. As automictic thelytokous parthenogenesis increases homozygosity over time (Pearcy, Hardy, & Aron, 2006; Pearcy, Hardy, & Aron, 2011), a decrease in observed heterozygosity and increase in relatedness should be present in the parthenogenetically produced queens when compared against sexually produced workers. For these and all further comparative analyses, figures were generated using the ggstatsplot R package (Patil, 2021).

### Chemical analyses

Chemical differentiation between nests was determined by analyzing eight randomly chosen workers per nest using GC–MS. Individual ants were knock-downed for one minute at −20 °C and extracted in 200µL hexane for five minutes with intermittent gentle mixing. Extracts were evaporated under a stream of high-purity nitrogen, redissolved in 35μL of hexane and transferred to a 100μL insert in a 1.5mL autoinjection vial. A volume of 2μl was injected in splitless mode using a 7693B Agilent autosampler into a HP-5MS UI column (30 m × 0.250 mm internal diameter × 0.25 μm film thickness; Agilent) with ultrahigh-purity helium as carrier gas (0.75 mL/min constant flow rate). The column was held at 50°C for 1 minute, increased to 320°C at 10°C/min, and held at 320°C for the last 10 minutes. The overall chemical profile of each individual was investigated by calculating the relative abundance of each compound. All compounds occurring in at least 10 samples were used to calculate the chemical profile of individuals, but we did not aim at identifying the different chemical compounds. The chemical profile of individuals were compared between nests, localities and populations.

We performed a PCA using ade4 (Dray & Dufour, 2007; R Core Team, 2020) in order to visualize the variation within and between nests. Additionally, we estimated the pairwise CHC differentiation between each nest through the calculation of the Euclidean distance between nest centroids. We then assessed the level of CHC variation within each nest by calculating the average Euclidean distance between each of the eight workers and the centroid of the nest. The between nest and within nest calculations were performed on the first two PC’s from the PCA. We first tested whether the level of CHC variation within nests differs between native and urban environments, as well as between monogyne and polygyne nests. Finally, we tested whether this level of CHC variation within a nest increases with genetic diversity (using expected heterozygosity as a proxy), and whether the level of CHC differentiation between nests increases with genetic differentiation or geographic distance, with significance determined using Student’s *t*-distribution for Pearson’s correlation coefficient.

### Behavioral assays

Aggression assays were performed by randomly selecting a single worker from two distinct nests and placing them together in a five-centimeter diameter petri dish for five minutes. The sides of the petri dish were coated with Fluon® to prevent the ants from escaping, and the bottom of the petri dish was covered with filter paper that was changed between trials to prevent odor transfer between trials. The subsequent behavioral interactions were scored on a four-level scale of escalating aggression (Suarez, Tsutsui, Holway, & Case, 1999): 1) touch (contacts that included prolonged antennation), 2) avoid (contacts that resulted in one or both ants quickly retreating in opposite directions), 3) aggression (lunging, biting, and pulling legs or antennae), or 4) fight (prolonged aggression between individuals). For each trial, the highest level of aggression was recorded, with the mean of 10 trials for each nest pairing used to calculate an average aggression score between nests. Twenty-five nest combinations were tested in Arkansas, California and Indiana (10 urban-urban, 10 natural-natural and 5 urban-natural) and 10 were tested in Colorado (10 natural-natural; no urban colonies found) for a total of 850 trials (85 nest combinations x 10 trials). Nests were matched across a short (minimum 0.001 kilometers) to long-distance (maximum 26 kilometers) gradient to identify whether geographic distance influenced aggression between nests. Similarly, aggression was compared against both genetic and chemical differentiation. The significance of all three relationships was evaluated using Student’s *t*-distribution for Pearson’s correlation coefficient. Finally, aggression levels among and between urban and natural nests were compared using Kruskal-Wallis tests, with Dunn’s test utilized to elucidate significant pairwise relationships and *p*-values adjusted by the Holm method to account for multiple comparisons.

## RESULTS

### Population and colony structure analyses

#### Mitochondrial dataset

A total of 68 mitochondrial haplotypes were identified, of which 36 were shared between individuals. The mean genetic distance between localities was 0.079, while it was only 0.039 within localities (Table S3). Mitochondrial haplotypes were rarely shared between localities (*i*.*e*., a single one shared between Arkansas and Indiana). Yet, the topology of the tree did not align completely with geography and therefore did not concur entirely with the four geographic clades previously uncovered by Menke et al. (2010) (Figure 1b; the mitochondrial network is provided in Figure S1). Notably, a substantial portion of the eastern US samples (*i*.*e*., Indiana and Arkansas) intermix with one another and appear on three distinct branches of the tree. Additionally, two samples from California were located nearest to the two basal eastern US branches, while samples from Colorado were split across two clades.

Our results further confirmed the finding of Menke et al. (2010) that mitochondrial haplotypes were commonly shared between monogyne and polygyne social structures (n = 12 – dispersed across all localities). However, haplotypes were rarely shared between natural and urban habitats, as only a single haplotype was shared between the two (in Indiana; Figures 1b and S1), suggesting that little genetic exchange occurs between habitats. Finally, the presence of multiple haplotypes within a nest was rare (n = 7 nests – only in the eastern US), suggesting polygyne colonies primarily develop via the association of related queens.

#### Microsatellite dataset

The 12 microsatellite markers used in this study contained an average of 12.6 alleles (range = 3 – 49; Table S4). When split by habitat, the natural and urban datasets contained an average of 9.8 (range = 3 – 38) and 9.2 (range = 3 – 37) alleles, respectively (Table S4). Therefore, the allelic diversity was not significantly different between natural and urban habitats (Mann-Whitney *U* = 4.31, *p* = 0.907). Furthermore, the allelic diversity was not significantly different between any of the four localities (Kruskal-Wallis *H* = 1.72, *p* = 0.633).

In the overall dataset, the AMOVA analysis revealed slight genetic diversity partitioned between localities (10.6%), with more substantial levels partitioned between habitats within localities (23.9%), between nests within habitats (27.2%), and within individuals themselves (45.5%; Table S5). The difference between localities is mostly driven by a clear separation of the eastern US samples (*i*.*e*., Indiana and Arkansas) from the western localities (*i*.*e*., Colorado and California) observed at *K* = 2 (Figure 1c, d). Consistently, Mantel tests identified significant isolation-by-distance when analyzing localities as a whole (Figure S2), contrasting the results obtained by the mitochondrial marker, where eastern and western populations did not segregate into two clearly distinct clades. Interestingly, at *K* = 4, the eastern US samples grouped by habitat rather than by locality, despite being geographically distant (Figure 1d). To a lesser extent, a similar pattern can be seen in the western localities, as the natural habitats of California and Colorado mostly grouped together (Figure 1d). At *K* = 7, both the localities and the habitats within localities clustered independently (Figure 1d), highlighting that the overall distribution of genetic variability is strongly influenced by both geographic distance and habitat.

Within localities, strong differentiation (*i*.*e*., high *F*_ST_) was found between almost every nest (Table 1, Figure S3). Similarly, *G* tests revealed that most nests represented a single genetic entity. Of the nest pairs that could not be differentiated, 11 were in the urban environment and two were in the natural environment (Supplementary Data). STRUCTURE analyses using the method of Puechmaille (2016) (*i*.*e*., finer partitioning) produced corroborating results, with best *K* mostly segregating each nest as its own genetic cluster (Figure 2). However, two trios of geographically adjacent nests were not genetically different from each other in urban habitats within California (nests 22-24) and Arkansas (nests 23-24, 29; Figures 1a, 2b, c). These two trios of nests also clustered together when urban habitats were analyzed separately from the natural habitat (Figure S4).

**Table 1:** The number and average number of alleles, observed (*H*_O_) and expected (*H*_E_) heterozygosity, inbreeding coefficient (*F*_IS_) and fixation index (*F*_ST_) for each locality and habitat across the 12 microsatellite loci. These statistics can be found for each microsatellite marker in Table S2.

| Location | Alleles | Average | $H_o$ | $H_e$ | $F_{IS}$ | $F_{ST}$ |
| --- | --- | --- | --- | --- | --- | --- |
| *Natural* |  |  |  |  |  |  |
| Indiana | 69 | 5.75 | 0.367 | 0.273 | -0.344 | 0.375 |
| Arkansas | 53 | 4.42 | 0.309 | 0.240 | -0.291 | 0.308 |
| California | 51 | 4.25 | 0.371 | 0.240 | -0.542 | 0.519 |
| Colorado | 53 | 4.42 | 0.310 | 0.228 | -0.360 | 0.391 |
| Overall | 117 | 9.75 | 0.339 | 0.245 | -0.384 | 0.591 |
| *Urban* |  |  |  |  |  |  |
| Indiana | 64 | 5.33 | 0.363 | 0.329 | -0.101 | 0.370 |
| Arkansas | 57 | 4.75 | 0.306 | 0.316 | 0.031 | 0.363 |
| California | 75 | 6.25 | 0.233 | 0.240 | 0.033 | 0.568 |
| Overall | 110 | 9.17 | 0.299 | 0.295 | -0.015 | 0.555 |
| All | 144 | 12.00 | 0.322 | 0.266 | -0.211 | 0.601 |

**Figure 2:**
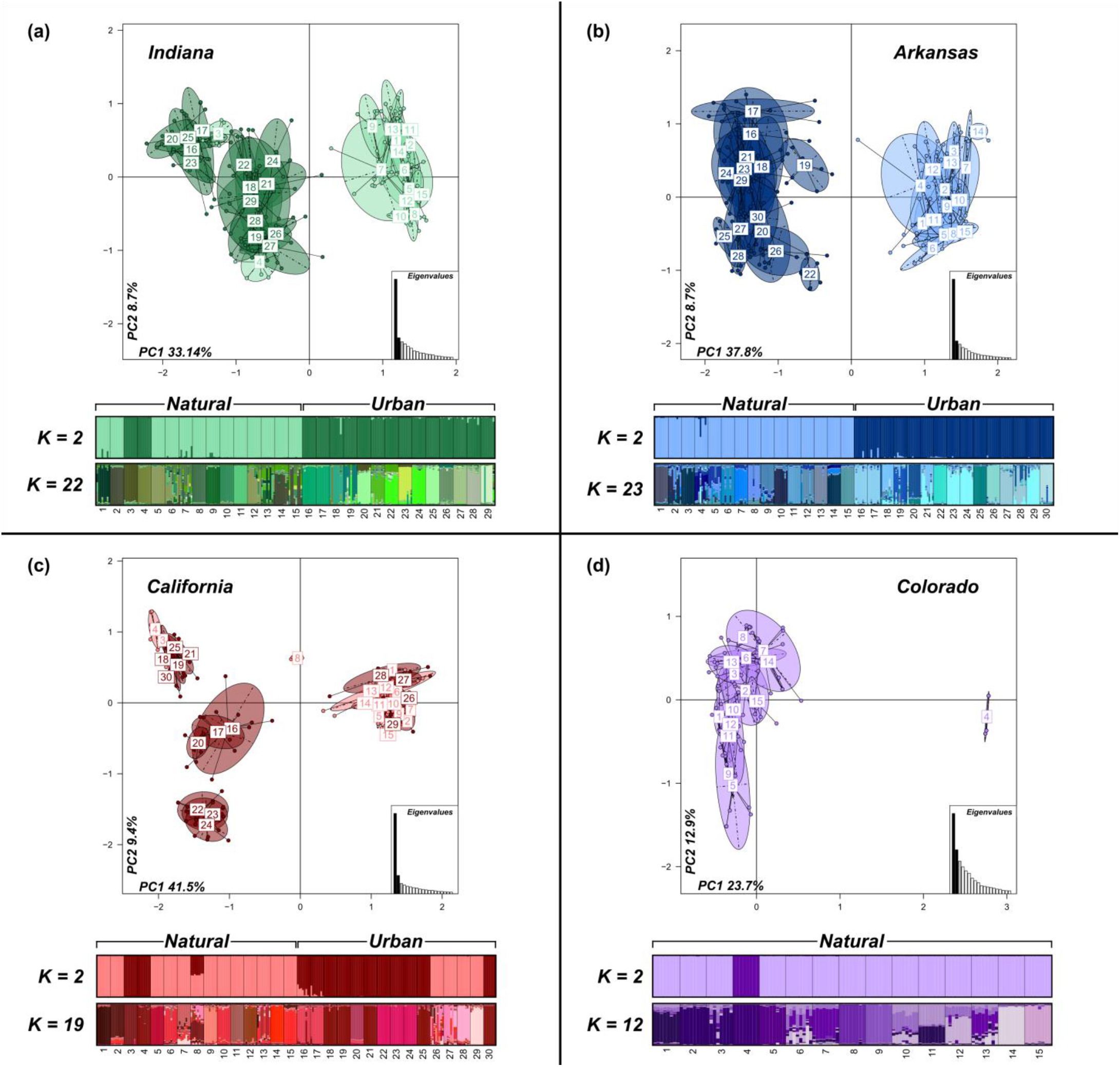
Population structure of *T. sessile* across the United States. Clustering of nests in **(a)** Indiana, **(b)** Arkansas, **(c)** California and **(d)** Colorado using a PCA and STRUCTURE on the microsatellite markers. For each locality, the light-shaded and dark-shaded ellipses in the PCA represent natural nests and urban nests, respectively. Additionally, two runs of STRUCTURE are shown for each locality, which correspond to best *K* (*i*.*e*., genetic clusters) as inferred by two different methods (Evanno above and Puechmaille below).

Remarkably, the Evanno et al. (2005) method (*i*.*e*., primary partitioning) consistently depicted clear separation between urban and natural habitats (Figure 2). *K* = 2 best explained the structure in the data for each locality and mostly segregated urban and natural colonies into two distinct clusters (Figure 2). This strong dichotomy between urban and natural habitats was also highlighted using PCAs within each locality (Figure 2). No isolation-by-distance was found when analyzing each habitat separately within localities (all *p* > 0.05; Figure S4); however, isolation-by-distance was significant when comparisons between habitats within localities were considered (*p* = 0.001, 0.001 and 0.016 for Indiana, Arkansas and California respectively; Figure 3a). Indeed, the inter-habitat genetic differentiation between nests was always higher than the differentiation between nests within a habitat (Figure 3b). However, the genetic differentiation between nests mostly did not differ across the two habitats (Figure 3b). AMOVAs for each locality were similar to the overall dataset, with most genetic diversity partitioned within individuals (avg. = 50%), and substantial amounts partitioned between habitats (avg. = 26%) and nests within each habitat (avg. = 31%; Table S6). Taken together, these results suggest that most nests sampled across the four localities represent distinct colonies. They also highlight the substantial differentiation between urban and natural populations, and support the continent-wide observation that colonies of *T. sessile* grouped by habitat rather than by locality within the eastern and western populations.

**Figure 3:**
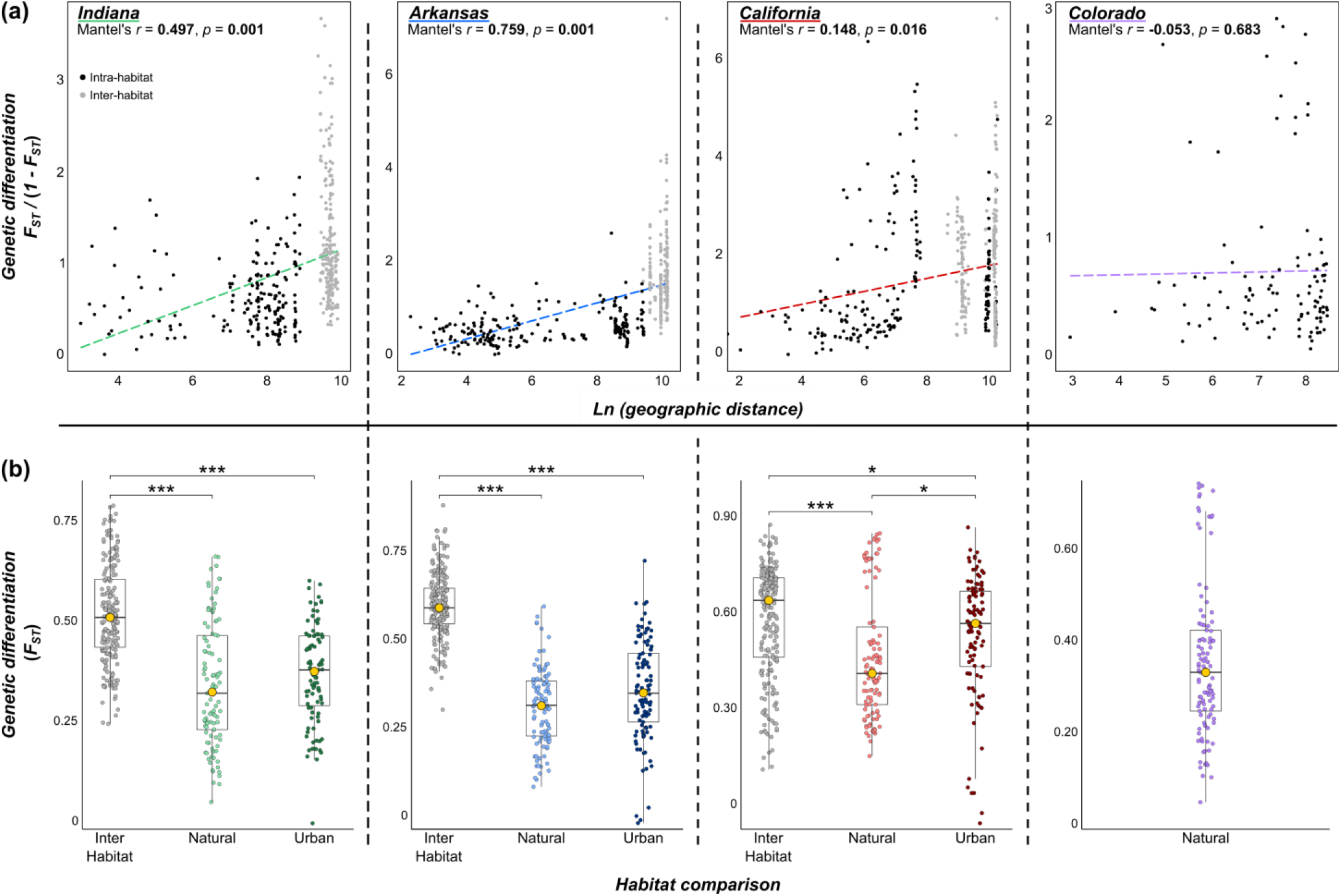
**(a)** Isolation-by-distance plots for each locality. Habitat specific (for Indiana, Arkansas and California) isolation-by-distance plots, as well as an overall isolation-by-distance plot, are available in the Supplementary Material. **(b)** Comparisons of genetic differentiation (*F*_ST_) between each pair of nests, both between and within habitats. Each gold dot on a boxplot represents the mean of the group, and only significant pairwise comparisons are shown (as determined by Dunn’s test with *p*-values adjusted according to the Holm method; \**p* < 0.05, \*\**p* < 0.01, \*\*\**p* < 0.001). Note that Colorado contains only intra-habitat comparisons for both **(a)** and **(b)**, as nests were only found in natural habitats.

### Breeding structure analyses

Overall, the urban environment contained significantly more polygyne nests (67%) than the natural environment (38%, p = 0.002; Figure 4a). Although both social structures were found in both habitats, the number of queens collected per nest was significantly higher in urban habitats (mean ± SD = 13.00 ± 15.70, up to 62) than natural habitats (mean ± SD = 2.61 ± 3.76, up to 21; Figure 4a). This pattern was found for the overall dataset (p < 0.001), as well as for each locality separately (despite being non-significant for Indiana, p = 0.117).

**Figure 4:**
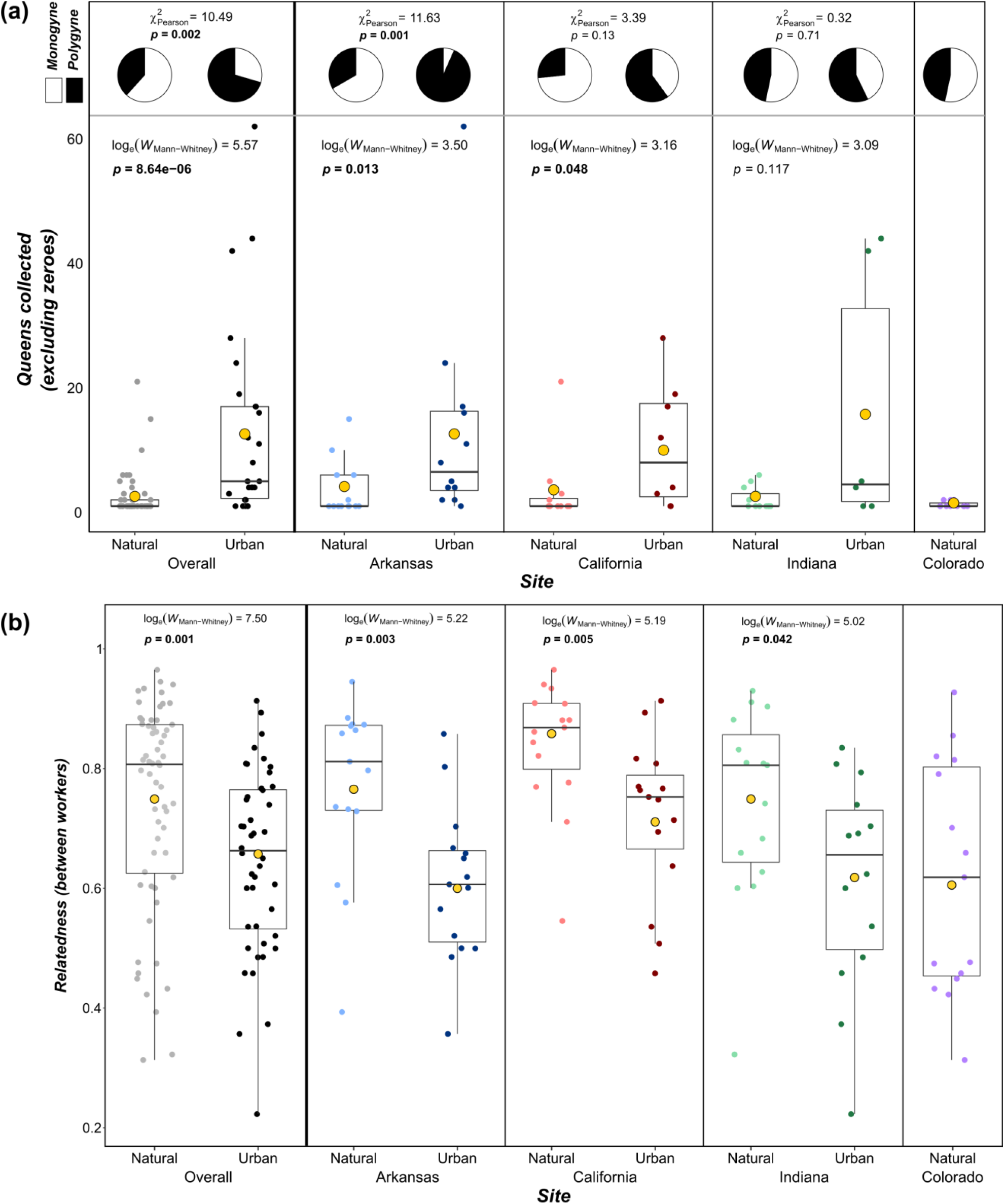
Breeding structure of *T. sessile* overall and across the four localities. **(a)** The percentage of monogyne and polygyne nests in natural and urban habitats, as well as the number of queens collected in nests where at least one queen was found. **(b)** The average relatedness among workers in urban and natural habitats. For both **(a)** and **(b)**, each smaller dot represents a nest, and each gold dot denotes the mean of the habitat.

Accordingly, the coefficient of relatedness between workers was significantly higher in the natural environment (R_w-w_ = 0.74) than in the urban environment (R_w-w_ = 0.65). This association was found significant for the overall dataset (*p* < 0.001), as well as for each locality separately (all *p* < 0.05; Figure 4b). However, the relatedness among workers was surprisingly high considering the number of queens present in each nest. This is especially true for urban nests (mean R_w-w_ = 0.61, 0.72 and 0.61 for the urban habitats of Arkansas, California and Indiana, respectively), as they usually contained a high number of queens. No values of relatedness were found close to zero (lowest value was 0.22), which is expected under a random association of a high number of queens or with the free movement of individuals among nests across the population.

Although natural nests were significantly more outbred (*i*.*e*., lower negative *F*_IS_) than their urban counterparts (*p* < 0.001; Table 1), the high relatedness within urban nests does not appear to stem from inbreeding (mean *F*_IS_ = −0.015). This suggests that queens do not exclusively participate in intranidal mating, although some level is likely considering the high relatedness values (and has been thought to occur in *T. sessile* – see Kannowski (1959)). Finally, the relatedness among queens within nests was also high (R_Q-Q_ = 0.68 and 0.66, for natural and urban polygyne nests, respectively), an uncommon finding for a polygynous ant and indicative of daughter queens being retained within their natal nest. However, this relatedness was not significantly higher (Figure S5) and the observed heterozygosity not significantly lower (Figure S6) when compared to the worker caste in any of the localities, suggesting that new queens are not produced asexually.

### Chemical analyses

Population clustering based on the cuticular hydrocarbons (CHCs) yielded similar results to those of the genetic analyses. At the overall scale, substantial chemical differentiation was found between localities, with the eastern US, California, and Colorado samples appearing distinct from one another (Figures S3 and S7). Consequently, CHC differentiation was significantly positively correlated with both geographic distance and genetic differentiation at the overall level (Figure S7). The Colorado samples were completely separate from the California samples, despite appearing genetically similar to samples from the California natural habitat. Interestingly, the urban and natural habitats from Arkansas, as well as the urban habitat from Indiana, clustered together chemically (Figures S3 and S7).

The chemical segregation of nests between natural and urban habitats became clearer when clustering analyses were performed at the locality level (Figure 5). Like the genetic differentiation, the chemical differentiation was mostly higher between nests from distinct habitats than between nests from the same habitat within a given locality (Figure 5). Consequently, the chemical differentiation between nests is associated within both genetic differentiation and geographic distance within most localities (Figure S8). Interestingly, nests 3 and 4 from the natural habitat of Indiana clustered with the urban habitat both genetically (Figure 2a) and chemically (Figure 5a). Overall, these findings suggest that chemical differentiation is influenced by both genetic and environmental factors, and therefore by the clear effect of habitat on the genetic differentiation mentioned above.

**Figure 5:**
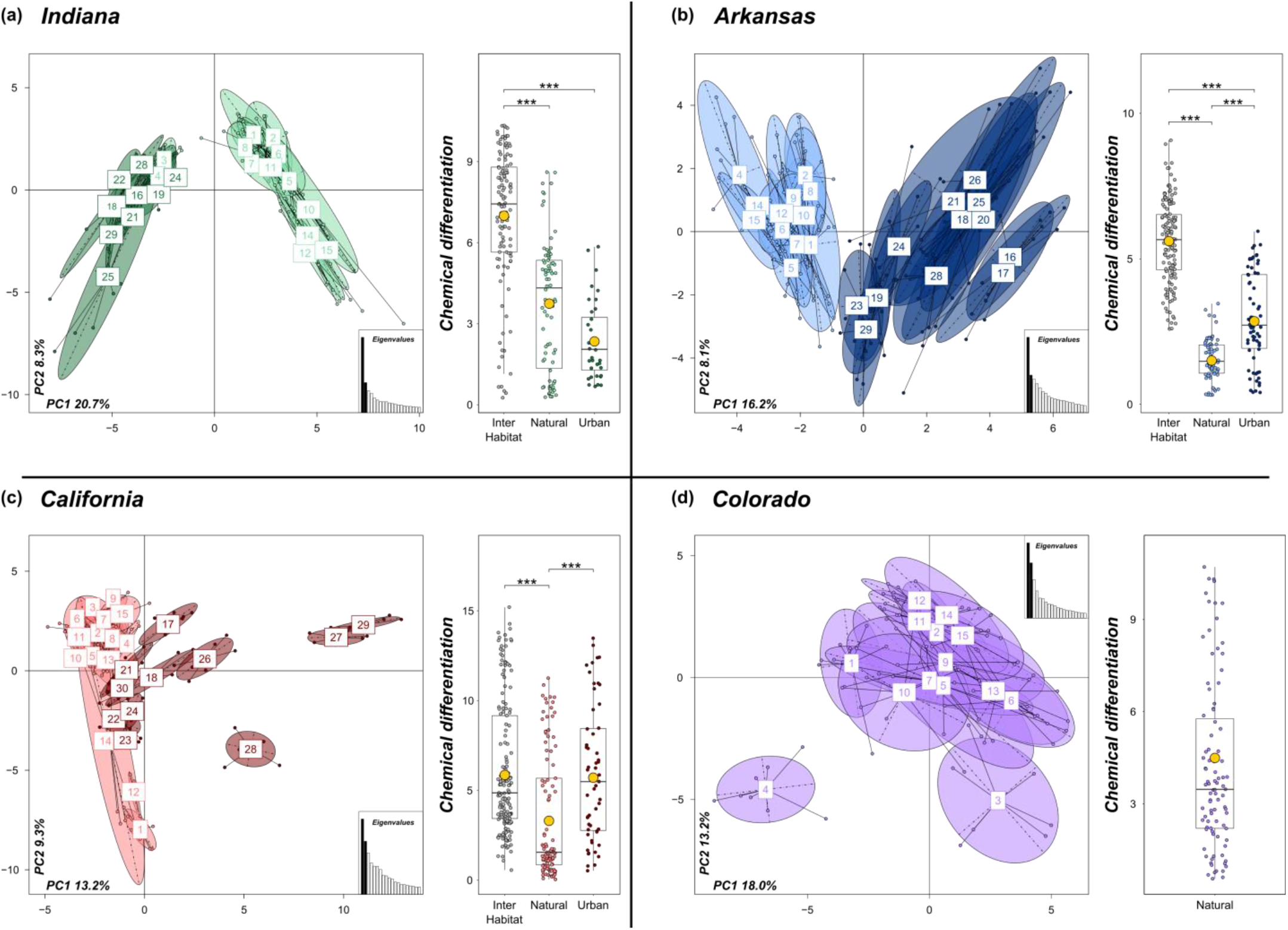
Clustering of nests in **(a)** Indiana, **(b)** Arkansas, **(c)** California and **(d)** Colorado using a PCA on each nest’s CHC profile. For each locality, the light-shaded and dark-shaded ellipses in the PCA represent natural nests and urban nests, respectively. Additionally, the boxplots illustrate the CHC differentiation of nests among and between habitats. Each smaller dot on the boxplots represents the difference between a pair of nests, while each gold dot denotes the mean of the group. Only significant pairwise comparisons are shown (as determined by Dunn’s test with *p*-values adjusted according to the Holm method; \**p* < 0.05, \*\**p* < 0.01, \*\*\**p* < 0.001). Clustering of nests based on CHC profiles in the overall population can be found in Figure S7.

Interestingly, the within-nest CHC variation was not significantly different between monogyne and polygyne nests in any locality or for the overall dataset. However, the within-nest variation was significantly different between natural and urban nests, with natural nests having increased variation at the overall level; a similar, but not significant, pattern was also observed within each locality (Figure S9). No significant correlations between CHC variation and genetic diversity were found within any locality (Figure S8).

### Behavioral assays

Aggression assays further demonstrated that most nests appear to be distinct colonies. The overwhelming majority of pairings obtained aggression scores of 3 or 4, with significant differences between groups mainly driven by slight fluctuations in avoidance/aggressive behaviors (Figure 6). The notable exception to these aggressive behaviors occurred in the California urban plot, where a more even distribution of aggression scores resulted from the lack of aggression between the trio of nests collected in San Francisco, California (nests 22-24; Figure 1a and 6). A similar trio of nests without aggression were also collected from the urban environment in Little Rock, Arkansas (nests 23-24, 29; see Figure 1a). Given that the genetic analyses above also support that the trio of nests in both cities comprise a single genetic entity, these two sites potentially represent supercolonies, albeit geographically limited (approximately 7,500 and 3,600 m^2^ in California and Arkansas, respectively).

**Figure 6:**
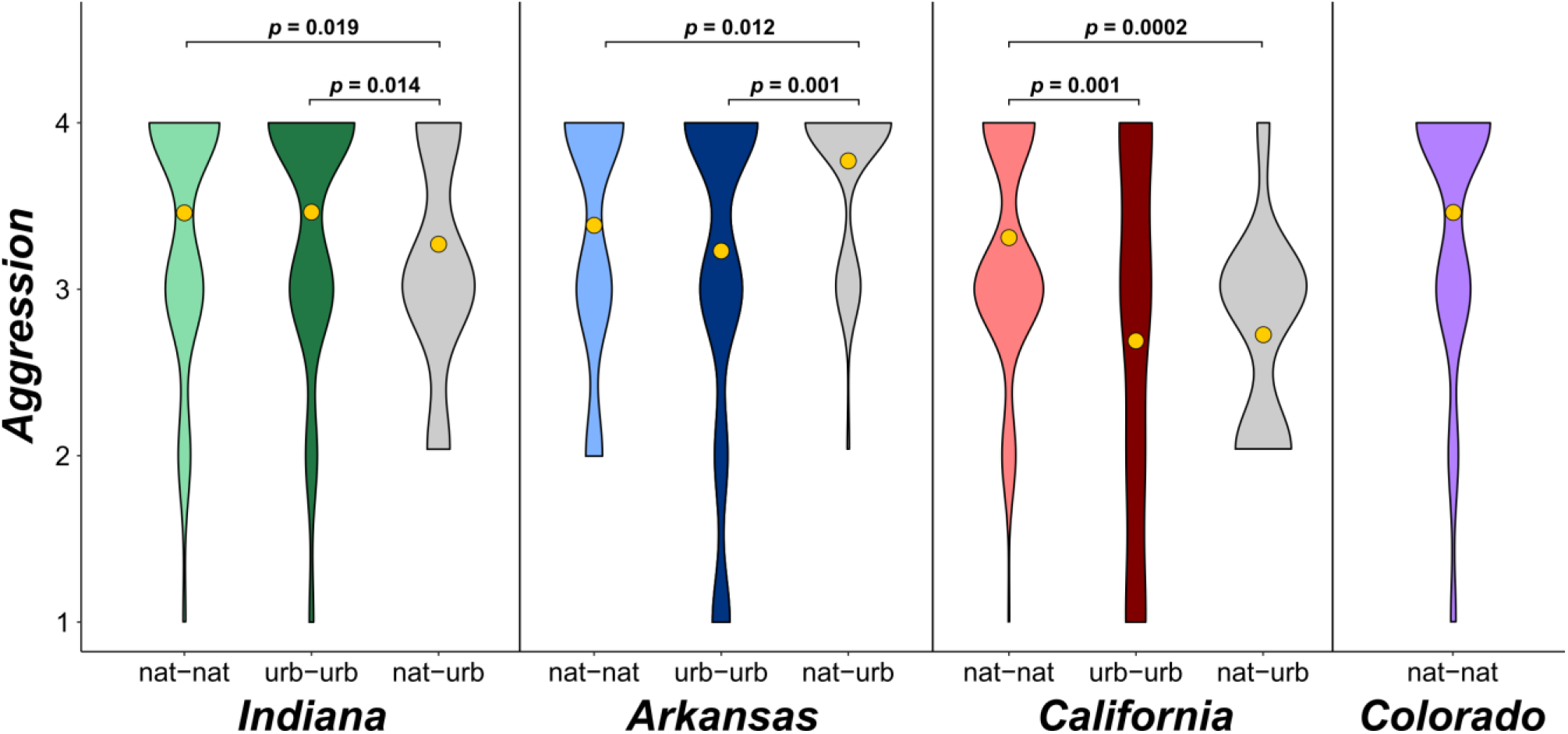
Violin plots of aggression between nests, with aggression compared within and among habitats for each locality (except Colorado). The gold dot on each violin represents the mean, and only significant pairwise comparisons are shown (as determined by Dunn’s test with *p*-values adjusted according to the Holm method).

The correlation analyses revealed little link between aggression and geographic distance, genetic differentiation or chemical differentiation (Figure S10), most likely driven by the high aggression that was observed in most aggression assays. Again, the notable exceptions here were the urban San Francisco and Little Rock sites mentioned above, as low genetic differentiation and aggression between some nests at these sites led to a significant positive correlation between genetic differentiation and aggression (Figure S10).

## DISCUSSION

Our extensive phylogenetic, chemical and behavioral study revealed several insights into the colony and breeding structure of *T. sessile* across the United States. First, we confirmed that the social structure of a colony appears to be a plastic trait found in both natural and urban habitats; however, we did find monogyny prevalent in natural habitats and polygyny largely associated with urban habitats. Furthermore, the extent of polygyny within nests was far greater in the urban habitat; however, differentiation was present between almost every nest in each locality and habitat. This finding, supported by our chemical and behavioral analyses, suggests that urban colonies mostly do not consist of large supercolonies. Yet, two trios of nests in two separate urban habitats lacked genetic, chemical and behavioral differentiation. As no such polydomous colonies were found in any natural habitat, this suggests only the urban environment harbors the necessary conditions for supercolony formation in *T. sessile*. Interestingly, our results uncovered clear genetic and chemical differentiation between natural and urban populations. This clear separation was observed within each locality, and also at a large geographic scale as nests clustered by habitat rather than by locality within the eastern and western clades. These findings suggest that urbanization acts as a strong barrier to gene flow in this species, with the heterogeneity of suitable habitat within cities potentially limiting immigration and emigration. Additionally, the clustering of nests by habitat rather than by locality may denote a strong signature of selection for the urban environment, with specific genotypes most likely found within cities. Overall, these results provide further support for urbanization as an intense driver of evolution.

Urbanization is a relatively recent byproduct of the Anthropocene (Lewis & Maslin, 2015); however, approximately 3% of Earth’s land surface is now urban and the rate of urbanization is accelerating (Liu, He, Zhou, & Wu, 2014). Urban environments are recognized as hotspots for microevolution due to the novel selection pressures arising from the changes of local biotic and abiotic interactions (Alberti, 2015; Johnson & Munshi-South, 2017). These changes include modifications to landscape composition (*e*.*g*., loss of suitable patches, homogenization and connectivity) (Groffman et al., 2014; McKinney, 2006), natural processes (*e*.*g*., soil pollution and nutrient cycling) (Isaksson, 2015) and ecological interactions (*e*.*g*., competition, predation and pathogens) (Rivkin et al., 2019). Modified ecological interactions within the urban environment affect the composition of communities, and have been shown to promote the abundance of arthropods (Faeth, Bang, & Saari, 2011), including ants (Vonshak & Gordon, 2015). Urban habitats may have allowed for *T. sessile* to achieve larger colony sizes than their natural conspecifics. For example, the abundance of traditionally limited resources may have stabilized, or even increased, colony productivity within urban habitats (Faeth, Warren, Shochat, & Marussich, 2005; Shochat, Warren, Faeth, McIntyre, & Hope, 2006). Additionally, increases in temperature in the winter (Arnfield, 2003; Oke, 1982) may extend the foraging and worker production season for urban colonies (Shochat et al., 2010; Shochat et al., 2006). Furthermore, fluctuations between seasons are lessened and extreme climate events buffered, which have been documented to prolong urban breeding seasons (Lowry, Lill, & Wong, 2013; Shochat et al., 2006). Therefore, these stable and highly productive environments may be providing favorable breeding conditions throughout the majority of the year, possibly contributing to the large colony sizes *T. sessile* attains in cities.

Nevertheless, although urbanization may reduce resource variation through time, resource variation in space may be intensified compared to natural habitats. Urban landscapes are often associated with a patchy distribution of resources (Cadenasso, Pickett, & Schwarz, 2007), which might have profound consequences on the evolution of dispersal traits in urban populations. Generally, costs of dispersal (*e*.*g*., loss of propagules, energetic demands) represent strong selective forces against dispersal (Bonte et al., 2012; Bowler & Benton, 2005). This is exemplified by the reduction in dispersal capability of island bird populations because a lack of predators nullifies the benefit flight offers as a predator-avoidance technique (Carlquist, 1966a; McNab, 1994; Wright, Steadman, & Witt, 2016), as well as in island plant and insect populations due to the high risk of landing in the ocean (Carlquist, 1966a, 1966b; Cody & Overton, 1996; Roff, 1990). Similarly, the fragmented habitats of urban environments may increase the failure rate of dispersers, and therefore select against dispersal. For example, urban populations of the pavement weed *Crepis sancta* produce a higher proportion of non-dispersing plants compared to their unfragmented natural populations (Cheptou, Carrue, Rouifed, & Cantarel, 2008). In ants, polygyny is usually selected for under conditions which diminish success rates of dispersal by young queens, such as ecological constraints on independent colony foundation (Keller, 1995). In a comparative study, Bourke and Heinze (1994) elucidated nest-site limitation, cold climate and habitat patchiness as factors that associate with polygyny in ants. Therefore, increased nest-site limitation due to the inherit patchiness within urban environments may be selecting for reduced dispersal of new queens, either through retention of new queens in their natal colony or dependent colony foundation (*i*.*e*., budding).

Notably, our results revealed surprisingly high levels of relatedness within urban colonies, including highly polygyne ones, suggesting queens are retained during colony growth. The retention of daughter queens may prevent the drastic loss of relatedness that the integration of new unrelated queens would otherwise cause, as well as prevent colony boundary collapse through a subsequent loss of non-nestmate discrimination. Interestingly, the development of large polydomous colonies observed in two urban localities was also not associated with a loss of relatedness among nestmate workers when compared to the genetic background of the locality. This finding supports the hypothesis that large polygyne/polydomous colonies arise through the extreme growth of a single colony (Helanterä, Strassmann, Carrillo, & Queller, 2009), likely through a combination of retaining daughter queens and the dependent colony foundation mentioned above. This hypothesis of supercolony development in invasive species supposes that polygyny and polydomy are pre-existing traits of the species within its native range (*e*.*g*., Fournier et al. (2012); Pedersen et al. (2006)); indeed, we found that these social traits were plastic in the natural habitat of *T. sessile*. This supercolony formation pathway allows for the coexistence of several competitive supercolonies within a given introduced/urban locality, a pattern found in many different invasive ant species (Abbott, 2005; Espadaler, Tartally, Schultz, Seifert, & Nagy, 2007; Giraud et al., 2002). Our results contrast with the two other hypotheses proposed to explain supercolony development in invasive ants. These hypotheses suggest that supercolonies arise from (i) a loss of nestmate recognition through a loss of diversity at the recognition loci or (ii) the selection for reduced aggression within densely populated areas (Helanterä et al., 2009). These two scenarios would both be accompanied by a loss of relatedness and reduced aggression among *all* urban nests, which was not found in our study.

Interestingly, we did not find significant isolation-by-distance within any habitat, although it is expected under dependent colony foundation. Rather, strong genetic differentiation was observed between nests regardless of geographic distance. The lack of isolation-by-distance in the urban habitat has a couple of possible explanations. For one, urban populations of *T. sessile* may have retained their dispersal capability despite facing a potentially increased cost of dispersal in patchy urban environments because the benefits of dispersal (*e*.*g*., inbreeding avoidance) outweigh the costs (*e*.*g*., not finding a suitable nesting site) (Bowler & Benton, 2005). However, each urban population in the study was significantly more inbred than their respective natural population despite not experiencing a reduction in genetic diversity, suggesting a shift in dispersal strategy between the habitats. Therefore, an alternative explanation for the lack of isolation-by-distance considering the increased inbreeding in urban populations may be sex-biased dispersal. As reduced dispersal may increase rates of inbreeding, male-biased dispersal could be selected for as a mechanism to avoid sibmating (Bowler & Benton, 2005). Indeed, both simulation models (Henry, Coulon, & Travis, 2016; Perrin & Mazalov, 2000) and empirical studies (Gauffre, Petit, Brodier, Bretagnolle, & Cosson, 2009; Oklander, Kowalewski, & Corach, 2010; Stow, Sunnucks, Briscoe, & Gardner, 2001) have found that polygynous systems select for male-biased dispersal, especially in fragmented habitats (*e*.*g*., cities). As dependent colony foundation in ants often results in a pattern of isolation-by-distance (Schultner et al., 2016), males of *T. sessile* may disperse far from their natal nest, rendering urban habitat isolation-by-distance non-significant. Therefore, dependent colony foundation and male-biased dispersal within urban populations may explain the elevated level of inbreeding compared to natural populations, as well the strong genetic sub-structure between most nests.

On the other hand, isolation-by-distance was found to be significant when analyzing both habitats together within a locality, highlighting the stark differentiation present between natural and urban populations. The minimal amount of gene flow between the two habitats suggests some selective force acting against inter-habitat colonization, as well as implies selection for certain traits within each environment. A genetic signature of rapid anthropogenic evolution has been purported for many species (Hendry, Farrugia, & Kinnison, 2008), although a more recent review found a lack of conclusive evidence for many studies (Lambert, Brans, Des Roches, Donihue, & Diamond, 2020). Rates of phenotypic change have been shown to be elevated in urban systems (Alberti et al., 2017), which are potentially driven by changes in the underlying genotypes. Variation in ecological conditions can certainly drive evolutionary shifts in species’ life history traits, but whether phenotypic shifts are primarily the result of genetic adaptations to urban environments via selection or simply phenotypic plasticity is unclear (Palkovacs & Hendry, 2010). Such a phenotypic shift has occurred in urban populations of the acorn ant *Temnothorax curvispinosus*, with urban workers exhibiting higher heat tolerance and diminished cold tolerance compared to natural populations (Diamond, Chick, Perez, Strickler, & Martin, 2018; Perez, Chick, Strickler, Diamond, & Zhao, 2018). A similar divergence was found between field-born workers and workers raised for two generations in the lab, indicating that such differences between habitats is driven by evolutionary divergence through selection rather than simply plasticity (Martin, Chick, Yilmaz, & Diamond, 2019). Reciprocal transplant experiments corroborated these results, with *T. curvispinosus* colonies experiencing higher survival in natal habitats compared to novel habitats, and local colonies displaying higher survival rates than foreign colonies (Martin, Chick, Garvin, & Diamond, 2021).

Interestingly, the two localities in the eastern United States grouped genetically by habitat rather than by locality. While this result makes sense for natural populations given the few barriers to gene flow between these regions of the country, it runs counter to several recent studies in other species which found heightened differentiation among distinct urban populations (Björklund, Ruiz, & Senar, 2010; Jason, Zolnik, & Harris, 2016; Lourenço, Álvarez, Wang, & Velo-Antón, 2017). However, gene flow between isolated urban populations may be sustained through human-mediated dispersal (Crispo, Moore, Lee-Yaw, Gray, & Haller, 2011). Such a scenario has been coined the ‘*urban facilitation model*’, whereby human-facilitated gene flow reduces differentiation between urban populations and may actually increase urban genetic diversity through the introduction of novel alleles (Miles, Johnson, Dyer, & Verrelli, 2018; Miles, Rivkin, Johnson, Munshi-South, & Verrelli, 2019). Indeed, invasion rates of ants have been shown to strongly correlate with waves of human globalization (Bertelsmeier et al., 2017), and a recent study on the tiny acorn ant *Temnothorax nylanderi* identified no significant differentiation between populations in distinct cities across France (Khimoun et al., 2020), highlighting that ants may be prime candidates for human-mediated dispersal. Considering selective pressures likely differ between the urban and nonurban habitats, connections between urban sites may facilitate the evolution of an “urban ecotype” (Schapira & Boutsika, 2012; Yakub & Tiffin, 2017). A more in-depth sampling scheme across the eastern United States is needed to test for such a gene flow assisted convergence in *Tapinoma sessile*.

## CONCLUSION

Convergent selection between distinct urban populations has been found across wide variety of taxa (Johnson, Prashad, Lavoignat, & Saini, 2018; Reid et al., 2016; Theodorou, Baltz, Paxton, & Soro, 2021; Yakub & Tiffin, 2017), suggesting attributes of the anthropogenic environment impart a homogenous influence upon evolution. Our study not only demonstrated the repeated life history shifts between natural and urban populations of T. sessile, but also highlighted the presence of significant genetic differentiation between these populations. Therefore, future studies are called for to clarify the mechanisms influencing the evolutionary trajectory of this ant. For example, transect sampling to evaluate whether the reduction of gene flow along the natural-to-urban habitat gradient is progressive or abrupt. Additionally, a genomic approach could help identify consistent genomic regions under selection to the urban environment across different US cities. Overall, these results reinforce the need for multi-faceted approaches in identifying signatures of local adaptation (Rivkin et al., 2019) and thereby the potential drivers of selection within cities (Lambert et al., 2020), as well as underscore the urban landscape as a powerful evolutionary force.

## Acknowledgements

This work was supported by the Urban Entomology Endowment at Texas A&M University and the Dr. Austin Frishman Scholarship from Pi Chi Omega.

## Author contributions

A.J.B. and E.L.V. designed the study. E.L.V. and A.M.H. contributed new reagents. A.J.B. collected the samples and performed the analyses. A.J.B. and P.A.E. analyzed the data. A.J.B. and P.A.E. wrote the paper with contributions of E.L.V., A.M.H. and G.B.

## Data availability statement

The data reported in this study have been deposited in the Open Science Framework database, https://osf.io (DOI 10.17605/OSF.IO/URKQV). Additionally, the mitochondrial COI sequences generated have been deposited to NCBI GenBank (Accession MZ007346-MZ007490).

## References

Abbott, K. L. (2005). Supercolonies of the invasive yellow crazy ant, Anoplolepis gracilipes, on an oceanic island: Forager activity patterns, density and biomass. Insectes Sociaux, 52(3), 266–273. doi:10.1007/s00040-005-0800-6

Alberti, M. (2015). Eco-evolutionary dynamics in an urbanizing planet. Trends in Ecology & Evolution, 30(2), 114–126. doi:10.1016/j.tree.2014.11.007

Alberti, M., Correa, C., Marzluff, J. M., Hendry, A. P., Palkovacs, E. P., Go-tanda, K. M., … Zhou, Y. (2017). Global urban signatures of phenotypic change in animal and plant populations. Proceedings of the National Academy of Sciences, USA, 114(34), 8951. doi:10.1073/pnas.1606034114

Arnfield, A. J. (2003). Two decades of urban climate research: a review of turbulence, exchanges of energy and water, and the urban heat island. International Journal of Climatology, 23(1), 1–26. doi:https://doi.org/10.1002/joc.859

Banks, N. C., Paini, D. R., Bayliss, K. L., & Hodda, M. (2015). The role of global trade and transport network topology in the human-mediated dispersal of alien species. Ecology Letters, 18(2), 188–199. doi:10.1111/ele.12397

Berman, M., Austin, C. M., & Miller, A. D. (2014). Characterisation of the complete mitochondrial genome and 13 microsatellite loci through next-generation sequencing for the New Caledonian spider-ant Leptomyrmex pallens. Molecular Biology Reports, 41(3), 1179–1187. doi:10.1007/s11033-013-2657-5

Bertelsmeier, C., Ollier, S., Liebhold, A., & Keller, L. (2017). Recent human history governs global ant invasion dynamics. Nature Ecology & Evolution, 1(7), 0184. doi:10.1038/s41559-017-0184

Beye, M., Neumann, P., Chapuisat, M., Pamilo, P., & Moritz, R. F. A. (1998). Nestmate recognition and the genetic relatedness of nests in the ant Formica pratensis. Behavioral Ecology and Sociobiology, 43(1), 67–72. doi:10.1007/s002650050467

Björklund, M., Ruiz, I., & Senar, J. C. (2010). Genetic differentiation in the urban habitat: the great tits (Parus major) of the parks of Barcelona city. Biological Journal of the Linnean Society, 99(1), 9–19. doi:10.1111/j.1095-8312.2009.01335.x

Bonte, D., Van Dyck, H., Bullock, J. M., Coulon, A., Delgado, M., Gibbs, M., … Travis, J. M. J. (2012). Costs of dispersal. Biological Reviews, 87(2), 290–312. doi:https://doi.org/10.1111/j.1469-185X.2011.00201.x

Boomsma, J. J., Huszár, D. B., & Pedersen, J. S. (2014). The evolution of multiqueen breeding in eusocial lineages with permanent physically differentiated castes. Animal Behaviour, 92, 241–252. doi:https://doi.org/10.1016/j.anbehav.2014.03.005

Boulay, R., Arnan, X., Cerdá, X., & Retana, J. (2014). The ecological benefits of larger colony size may promote polygyny in ants. Journal of Evolutionary Biology, 27(12), 2856–2863. doi:doi:10.1111/jeb.12515

Bourke, A. F. G., & Heinze, J. (1994). The ecology of communal breeding: the case of multiple-queen Leptothoracine ants. Philosophical Transactions of the Royal Society of London. Series B: Biological Sciences, 345(1314), 359–372. doi:doi:10.1098/rstb.1994.0115

Boutin-Ganache, I., Raposo, M., Raymond, M., & Deschepper, C. F. (2001). M13-tailed primers improve the readability and usability of microsatellite analyses performed with two different allele-sizing methods. BioTechniques, 31(1).

Bowler, D. E., & Benton, T. G. (2005). Causes and consequences of animal dispersal strategies: relating individual behaviour to spatial dynamics. Biological Reviews, 80(2), 205–225. doi:https://doi.org/10.1017/S1464793104006645

Buczkowski, G. (2010). Extreme life history plasticity and the evolution of invasive characteristics in a native ant. Biological Invasions, 12(9), 3343–3349. doi:10.1007/s10530-010-9727-6

Buczkowski, G., & Bennett, G. (2008). Seasonal polydomy in a polygynous supercolony of the odorous house ant, Tapinoma sessile. Ecological Entomology, 33, 780–788. doi:10.1111/j.1365-2311.2008.01034.x

Buczkowski, G., & Bennett, G. (2009). Colony budding and its effects on food allocation in the highly polygynous ant, Monomorium pharaonis. Ethology, 115(11), 1091–1099. doi:https://doi.org/10.1111/j.1439-0310.2009.01698.x

Buczkowski, G., & Krushelnycky, P. (2011). The odorous house ant, Tapinoma sessile (Hymenoptera: Formicidae), as a new temperate-origin invader. Myrmecological News, 16, 61–66.

Butler, I. A., Siletti, K., Oxley, P. R., & Kronauer, D. J. C. (2014). Conserved Microsatellites in Ants Enable Population Genetic and Colony Pedigree Studies across a Wide Range of Species. PLoS One, 9(9), e107334. doi:10.1371/journal.pone.0107334

Cadenasso, M. L., Pickett, S. T. A., & Schwarz, K. (2007). Spatial heterogeneity in urban ecosystems: reconceptualizing land cover and a framework for classification. Frontiers in Ecology and the Environment, 5(2), 80–88. doi:https://doi.org/10.1890/1540-9295(2007)5[80:SHIUER]20.CO;2

Carlquist, S. (1966a). The biota of long-distance dispersal. I. principles of dispersal and evolution. The Quarterly Review of Biology, 41(3), 247–270. doi:10.1086/405054

Carlquist, S. (1966b). The biota of long-distance dispersal. II. Loss of dispersibility in Pacific Compositae. Evolution, 20(1), 30–48. doi:10.2307/2406147

Chapuisat, M., Bernasconi, C., Hoehn, S., & Reuter, M. (2005). Nestmate recognition in the unicolonial ant Formica paralugubris. Behavioral Ecology, 16(1), 15–19. doi:10.1093/beheco/arh128

Cheptou, P. O., Carrue, O., Rouifed, S., & Cantarel, A. (2008). Rapid evolution of seed dispersal in an urban environment in the weed Crepis sancta. Proceedings of the National Academy of Sciences, USA, 105(10), 3796. doi:10.1073/pnas.0708446105

Cody, M. L., & Overton, J. M. (1996). Short-term evolution of reduced dispersal in island plant populations. Journal of Ecology, 84(1), 53–61. doi:10.2307/2261699

Crispo, E., Moore, J.-S., Lee-Yaw, J. A., Gray, S. M., & Haller, B. C. (2011). Broken barriers: human-induced changes to gene flow and introgression in animals. BioEssays, 33(7), 508–518. doi:https://doi.org/10.1002/bies.201000154

Cronin, A. L., Molet, M., Doums, C., Monnin, T., & Peeters, C. (2013). Recurrent evolution of dependent colony foundation across eusocial insects. Annual Review of Entomology, 58(1), 37–55. doi:10.1146/annurev-ento-120811-153643

Diamond, S. E., Chick, L. D., Perez, A., Strickler, S. A., & Martin, R. A. (2018). Evolution of thermal tolerance and its fitness consequences: parallel and non-parallel responses to urban heat islands across three cities. Proceedings of the Royal Society B: Biological Sciences, 285(1882), 20180036. doi:doi:10.1098/rspb.2018.0036

Dray, S., & Dufour, A.-B. (2007). The ade4 package: implementing the duality diagram for ecologists. Journal of Statistical Software, 22(4), 1–20.

Du, Y., Song, X., Yu, H., & Lu, Z. (2019). Complete mitochondrial genome sequence of Tapinoma melanocephalum (Hymenoptera: Formicidae). Mitochondrial DNA Part B, 4(2), 3448–3449. doi:10.1080/23802359.2019.1674205

Dussutour, A., & Simpson, S. J. (2008). Description of a simple synthetic diet for studying nutritional responses in ants. Insectes Sociaux, 55(3), 329–333. doi:10.1007/s00040-008-1008-3

Ehrenfeld, J. G. (2010). Ecosystem Consequences of Biological Invasions. Annual Review of Ecology, Evolution, and Systematics, 41(1), 59–80. doi:10.1146/annurev-ecolsys-102209-144650

Espadaler, X., Tartally, A., Schultz, R., Seifert, B., & Nagy, C. (2007). Regional trends and preliminary results on the local expansion rate in the invasive garden ant, Lasius neglectus (Hymenoptera, Formicidae). Insectes Sociaux, 54(3), 293–301. doi:10.1007/s00040-007-0944-7

Evanno, G., Regnaut, S., & Goudet, J. (2005). Detecting the number of clusters of individuals using the software STRUCTURE: a simulation study. Molecular Ecology, 14(8), 2611–2620. doi:10.1111/j.1365-294X.2005.02553.x

Eyer, P.-A., McDowell, B., Johnson, L. N. L., Calcaterra, L. A., Fernandez, M. B., Shoemaker, D., Vargo, E. L. (2018). Supercolonial structure of invasive populations of the tawny crazy ant Nylanderia fulva in the US. BMC Evolutionary Biology, 18(1), 209. doi:10.1186/s12862-018-1336-5

Eyer, P.-A., & Vargo, E. L. (2021). Breeding structure and invasiveness in social insects. Current Opinion in Insect Science, 46, 24–30. doi:https://doi.org/10.1016/j.cois.2021.01.004

Faeth, S. H., Bang, C., & Saari, S. (2011). Urban biodiversity: patterns and mechanisms. Annals of the New York Academy of Sciences, 1223(1), 69–81.

Faeth, S. H., Warren, P. S., Shochat, E., & Marussich, W. A. (2005). Trophic dynamics in urban communities. BioScience, 55(5), 399–407. doi:10.1641/0006-3568(2005)055[0399:Tdiuc]20.Co;2

Falush, D., Stephens, M., & Pritchard, J. K. (2003). Inference of population structure using multilocus genotype data: linked loci and correlated allele frequencies. Genetics, 164(4), 1567.

Fournier, D., Estoup, A., Orivel, J., Foucaud, J., Jourdan, H., Breton, J. L., & Keller, L. (2005). Clonal reproduction by males and females in the little fire ant. Nature, 435(7046), 1230–1234. doi:10.1038/nature03705

Fournier, D., Tindo, M., Kenne, M., Mbenoun Masse, P. S., Van Bossche, V., De Coninck, E., Fenton, B. (2012). Genetic structure, nestmate recognition and behaviour of two cryptic species of the invasive big-headed ant Pheidole megacephala. PLoS ONE, 7(2), e31480.

Gauffre, B., Petit, E., Brodier, S., Bretagnolle, V., & Cosson, J. F. (2009). Sex-biased dispersal patterns depend on the spatial scale in a social rodent. Proceedings of the Royal Society B: Biological Sciences, 276(1672), 3487–3494. doi:doi:10.1098/rspb.2009.0881

Giraud, T., Pedersen, J. S., & Keller, L. (2002). Evolution of supercolonies: the Argentine ants of southern Europe. Proceedings of the National Academy of Sciences, USA, 99(9), 6075. doi:10.1073/pnas.092694199

Goudet, J. (2003). FSTAT (ver. 2.9.4), a program to estimate and test population genetics parameters. Available from http://www.unil.ch/izea/soft-wares/fstat.html Updated from Goudet [1995].

Groffman, P. M., Cavender-Bares, J., Bettez, N. D., Grove, J. M., Hall, S. J., Heffernan, J. B., Steele, M. K. (2014). Ecological homogenization of urban USA. Frontiers in Ecology and the Environment, 12(1), 74–81. doi:https://doi.org/10.1890/120374

Hajibabaei, M., Janzen, D. H., Burns, J. M., Hallwachs, W., & Hebert, P. D. N. (2006). DNA barcodes distinguish species of tropical Lepidoptera. Proceedings of the National Academy of Sciences of the United States of America, 103(4), 968. doi:10.1073/pnas.0510466103

Hebert, P. D. N., Penton, E. H., Burns, J. M., Janzen, D. H., & Hallwachs, W. (2004). Ten species in one: DNA barcoding reveals cryptic species in the neotropical skipper butterfly *Astraptes fulgerator*. Proceedings of the National Academy of Sciences of the United States of America, 101(41), 14812. doi:10.1073/pnas.0406166101

Helanterä, H., Strassmann, J. E., Carrillo, J., & Queller, D. C. (2009). Unicolonial ants: where do they come from, what are they and where are they going? Trends in Ecology & Evolution, 24(6), 341–349.

Hendry, A. P., Farrugia, T. J., & Kinnison, M. T. (2008). Human influences on rates of phenotypic change in wild animal populations. Molecular Ecology, 17(1), 20–29. doi:https://doi.org/10.1111/j.1365-294X.2007.03428.x

Henry, R. C., Coulon, A., & Travis, J. M. J. (2016). The evolution of male-biased dispersal under the joint selective forces of inbreeding load and demographic and environmental stochasticity. The American Naturalist, 188(4), 423–433. doi:10.1086/688170

Hölldobler, B., & Wilson, E. O. (1977). The number of queens: An important trait in ant evolution. Naturwissenschaften, 64(1), 8–15. doi:10.1007/bf00439886

Holm, S. (1979). A simple sequentially rejective multiple test procedure. Scandinavian Journal of Statistics, 6(2), 65–70.

Holway, D. A. (1999). Competitive mechanisms underlying the displacement of native ants by the invasive Argentine ant. Ecology, 80(1), 238–251. doi:https://doi.org/10.1890/0012-9658(1999)080[0238:CMUTDO]2.0.CO;2

Holway, D. A., Suarez, A. V., & Case, T. J. (1998). Loss of intraspecific aggression in the success of a widespread invasive social insect. Science, 282(5390), 949. doi:10.1126/science.282.5390.949

Isaksson, C. (2015). Urbanization, oxidative stress and inflammation: a question of evolving, acclimatizing or coping with urban environmental stress. Functional Ecology, 29(7), 913–923. doi:https://doi.org/10.1111/1365-2435.12477

Jason, M. S., Zolnik, C. P., & Harris, S. E. (2016). Population genomics of the Anthropocene: urbanization is negatively associated with genome-wide variation in white-footed mouse populations. Evolutionary Applications, 9(4), 546–564. doi:http://dx.doi.org/10.1111/eva.12357

Jeschke, J., & Wittenborn, D. (2011). Characteristics of exotic ants in North America. NeoBiota, 10, 47–64. doi:10.3897/neobiota.10.1047

Johnson, M. T. J., & Munshi-South, J. (2017). Evolution of life in urban environments. Science, 358(6363), eaam8327. doi:10.1126/science.aam8327

Johnson, M. T. J., Prashad, C. M., Lavoignat, M., & Saini, H. S. (2018). Contrasting the effects of natural selection, genetic drift and gene flow on urban evolution in white clover (Trifolium repens). Proceedings of the Royal Society B: Biological Sciences, 285(1883), 20181019. doi:doi:10.1098/rspb.2018.1019

Jombart, T. (2008). adegenet: a R package for the multivariate analysis of genetic markers. Bioinformatics, 24(11), 1403–1405.

Kamvar, Z. N., Tabima, J. F., & Grünwald, N. J. (2014). Poppr: an R package for genetic analysis of populations with clonal, partially clonal, and/or sexual reproduction. PeerJ, 2, e281. doi:10.7717/peerj.281

Kannowski, P. B. (1959). The flight activities and colony-founding behavior of bog ants in Southeastern Michigan. Insectes Sociaux, 6(2), 115–162. doi:10.1007/BF02225947

Katzerke, A., Neumann, P., Pirk, C. W. W., Bliss, P., & Moritz, R. F. A. (2006). Seasonal nestmate recognition in the ant Formica exsecta. Behavioral Ecology and Sociobiology, 61(1), 143–150. doi:10.1007/s00265-006-0245-6

Kearse, M., Moir, R., Wilson, A., Stones-Havas, S., Cheung, M., Sturrock, S., … Drummond, A. (2012). Geneious Basic: an integrated and extendable desktop software platform for the organization and analysis of sequence data. Bioinformatics, 28(12), 1647–1649. doi:10.1093/bioinformatics/bts199

Keller, L. (1991). Queen number, mode of colony founding, and queen reproductive success in ants (Hymenoptera Formicidae). Ethology Ecology & Evolution, 3(4), 307–316. doi:10.1080/08927014.1991.9525359

Keller, L. (1995). Social life: the paradox of multiple-queen colonies. Trends in Ecology & Evolution, 10(9), 355–360. doi:https://doi.org/10.1016/S0169-5347(00)89133-8

Khimoun, A., Doums, C., Molet, M., Kaufmann, B., Peronnet, R., Eyer, P. A., & Mona, S. (2020). Urbanization without isolation: the absence of genetic structure among cities and forests in the tiny acorn ant Temnothorax nylanderi. Biology Letters, 16(1), 20190741. doi:doi:10.1098/rsbl.2019.0741

Kimura, M. (1980). A simple method for estimating evolutionary rates of base substitutions through comparative studies of nucleotide sequences. Journal of Molecular Evolution, 16(2), 111–120. doi:10.1007/BF01731581

Kopelman, N. M., Mayzel, J., Jakobsson, M., Rosenberg, N. A., & Mayrose, I. (2015). Clumpak: a program for identifying clustering modes and packaging population structure inferences across K. Molecular Ecology Resources, 15(5), 1179–1191. doi:10.1111/1755-0998.12387

Krieger, M. J. B., & Keller, L. (1999). Low polymorphism at 19 microsatellite loci in a French population of Argentine ants (Linepithema humile). Molecular Ecology, 8, 1078–1080.

Kumar, S., Stecher, G., Li, M., Knyaz, C., & Tamura, K. (2018). MEGA X: Molecular evolutionary genetics analysis across computing platforms. Molecular Biology and Evolution, 35(6), 1547–1549. doi:10.1093/mol-bev/msy096

Lambert, M. R., Brans, K. I., Des Roches, S., Donihue, C. M., & Diamond, S. E. (2020). Adaptive evolution in cities: progress and misconceptions. Trends in Ecology & Evolution. doi:https://doi.org/10.1016/j.tree.2020.11.002

Leigh, J. W., & Bryant, D. (2015). POPART: full-feature software for haplotype network construction. Methods in Ecology and Evolution, 6(9), 1110–1116. doi:https://doi.org/10.1111/2041-210X.12410

Lester, P. J., & Gruber, M. A. M. (2016). Booms, busts and population collapses in invasive ants. Biological Invasions, 18(11), 3091–3101. doi:10.1007/s10530-016-1214-2

Lewis, S. L., & Maslin, M. A. (2015). Defining the Anthropocene. Nature, 519(7542), 171–180. doi:10.1038/nature14258

Li, Y.-L., & Liu, J.-X. (2018). StructureSelector: A web-based software to select and visualize the optimal number of clusters using multiple methods. Molecular Ecology Resources, 18(1), 176–177. doi:https://doi.org/10.1111/1755-0998.12719

Liu, Z., He, C., Zhou, Y., & Wu, J. (2014). How much of the world’s land has been urbanized, really? A hierarchical framework for avoiding confusion. Landscape Ecology, 29(5), 763–771. doi:10.1007/s10980-014-0034-y

Lockey, K. H. (1991). Insect hydrocarbon classes: Implications for chemotaxonomy. Insect Biochemistry, 21(1), 91–97. doi:https://doi.org/10.1016/0020-1790(91)90068-P

Lourenço, A., Álvarez, D., Wang, I. J., & Velo-Antón, G. (2017). Trapped within the city: integrating demography, time since isolation and population-specific traits to assess the genetic effects of urbanization. Molecular Ecology, 26(6), 1498–1514. doi:https://doi.org/10.1111/mec.14019

Lowry, H., Lill, A., & Wong, B. B. M. (2013). Behavioural responses of wildlife to urban environments. Biological Reviews, 88(3), 537–549. doi:https://doi.org/10.1111/brv.12012

Martin, R. A., Chick, L. D., Garvin, M. L., & Diamond, S. E. (2021). In a nutshell, a reciprocal transplant experiment reveals local adaptation and fitness trade-offs in response to urban evolution in an acorn-dwelling ant. Evolution, 75(4), 876–887. doi:https://doi.org/10.1111/evo.14191

Martin, R. A., Chick, L. D., Yilmaz, A. R., & Diamond, S. E. (2019). Evolution, not transgenerational plasticity, explains the adaptive divergence of acorn ant thermal tolerance across an urban–rural temperature cline. Evolutionary Applications, 12(8), 1678–1687. doi:10.1111/eva.12826

McGlynn, T. P. (1999). The worldwide transfer of ants: geographical distribution and ecological invasions. Journal of Biogeography, 26, 535–548.

McKinney, M. L. (2006). Urbanization as a major cause of biotic homogenization. Biological Conservation, 127(3), 247–260. doi:https://doi.org/10.1016/j.biocon.2005.09.005

McNab, B. K. (1994). Energy conservation and the evolution of flightlessness in birds. The American Naturalist, 144(4), 628–642.

Menke, S. B., Booth, W., Dunn, R. R., Schal, C., Vargo, E. L., & Silverman, J. (2010). Is it easy to be urban? Convergent success in urban habitats among lineages of a widespread native ant. PLOS One, 5(2), e9194. doi:10.1371/journal.pone.0009194

Meyerson, L. A., & Mooney, H. A. (2007). Invasive alien species in an era of globalization. Frontiers in Ecology and the Environment, 5(4), 199–208. doi:https://doi.org/10.1890/1540-9295(2007)5[199:IASIAE]20.CO;2

Miles, L. S., Johnson, J. C., Dyer, R. J., & Verrelli, B. C. (2018). Urbanization as a facilitator of gene flow in a human health pest. Molecular Ecology, 27(16), 3219–3230. doi:https://doi.org/10.1111/mec.14783

Miles, L. S., Rivkin, L. R., Johnson, M. T. J., Munshi-South, J., & Verrelli, B. C. (2019). Gene flow and genetic drift in urban environments. Molecular Ecology, 28(18), 4138–4151. doi:https://doi.org/10.1111/mec.15221

Nei, M. (1987). Molecular Evolutionary Genetics: Columbia University Press.

Oke, T. R. (1982). The energetic basis of the urban heat island. Quarterly Journal of the Royal Meteorological Society, 108(455), 1–24. doi:https://doi.org/10.1002/qj.49710845502

Oklander, L. I., Kowalewski, M. M., & Corach, D. (2010). Genetic consequences of habitat fragmentation in glack-and-gold howler (Alouatta caraya) populations from Northern Argentina. International Journal of Primatology, 31(5), 813–832. doi:http://dx.doi.org/10.1007/s10764-010-9430-6

Oksanen, J., Blanchet, F. G., Friendly, M., Kindt, R., Legendre, P., McGlinn, D., & Minchin, P. R. (2020). vegan: Community Ecology Package. https://CRAN.R-project.org/package=vegan.

Palkovacs, E. P., & Hendry, A. P. (2010). Eco-evolutionary dynamics: intertwining ecological and evolutionary processes in contemporary time. F1000 Biology Reports, 2, 1. doi:10.3410/B2-1

Patil, I. (2021). Visualizations with statistical details: The’ggstatsplot’approach. The Journal of Open Source Software, 6(61), 3167. doi:10.21105/joss.03167

Pearcy, M., Goodisman, M. A., & Keller, L. (2011). Sib mating without inbreeding in the longhorn crazy ant. Proceedings of the Royal Society B: Biological Sciences, 278(1718), 2677–2681. doi:10.1098/rspb.2010.2562

Pearcy, M., Hardy, O., & Aron, S. (2006). Thelytokous parthenogenesis and its consequences on inbreeding in an ant. Heredity, 96(5), 377–382. doi:10.1038/sj.hdy.6800813

Pearcy, M., Hardy, O. J., & Aron, S. (2011). Automictic parthenogenesis and rate of transition to homozygosity. Heredity, 107(2), 187–188. doi:10.1038/hdy.2010.172

Pedersen, J. S., Krieger, M. J. B., Vogel, V., Giraud, T., & Keller, L. (2006). Native supercolonies of unrelated individuals in the invasive Argentine ant. Evolution, 60(4), 782-791, 710.

Perez, A., Chick, L. D., Strickler, S. A., Diamond, S. E., & Zhao, C. (2018). Evolution of plasticity in the city: urban acorn ants can better tolerate more rapid increases in environmental temperature. Conservation Physiology, 6(1). doi:10.1093/conphys/coy030

Perrin, N., & Mazalov, V. (2000). Local competition, inbreeding, and the evolution of sex-biased dispersal. The American Naturalist, 155(1), 116–127. doi:10.1086/303296

Pirk, C. W. W., Neumann, P., Moritz, R. F. A., & Pamilo, P. (2001). Intranest relatedness and nestmate recognition in the meadow ant Formica pratensis (R.). Behavioral Ecology and Sociobiology, 49(5), 366–374. doi:10.1007/s002650000315

Pritchard, J. K., Stephens, M., & Donnelly, P. (2000). Inference of population structure using multilocus genotype data. Genetics, 155, 945–959.

Puechmaille, S. J. (2016). The program STRUCTURE does not reliably recover the correct population structure when sampling is uneven: subsampling and new estimators alleviate the problem. Molecular Ecology Resources, 16(3), 608–627. doi:https://doi.org/10.1111/1755-0998.12512

Queller, D. C., & Goodnight, K. F. (1989). Estimating relatedness using genetic markers. Evolution, 43(2), 258–275. doi:doi:10.1111/j.1558-5646.1989.tb04226.x

R Core Team. (2020). R: A language and environment for statistical computing. Vienna, Austria: R Foundation for Statistical Computing. Retrieved from https://www.R-project.org/

Rabeling, C., & Kronauer, D. J. C. (2013). Thelytokous parthenogenesis in eusocial Hymenoptera. Annual Review of Entomology, 58(1), 273–292. doi:10.1146/annurev-ento-120811-153710

Reid, N. M., Proestou, D. A., Clark, B. W., Warren, W. C., Colbourne, J. K., Shaw, J. R., … Whitehead, A. (2016). The genomic landscape of rapid repeated evolutionary adaptation to toxic pollution in wild fish. Science, 354(6317), 1305. doi:10.1126/science.aah4993

Rivkin, L. R., Santangelo, J. S., Alberti, M., Aronson, M. F. J., de Keyzer, C. W., Diamond, S. E., … Johnson, M. T. J. (2019). A roadmap for urban evolutionary ecology. Evolutionary Applications, 12(3), 384–398. doi:https://doi.org/10.1111/eva.12734

Roff, D. A. (1990). The evolution of flightlessness in insects. Ecological Monographs, 60(4), 389–421. doi:10.2307/1943013

Röhl, A., Bandelt, H. J., & Forster, P. (1999). Median-joining networks for inferring intraspecific phylogenies. Molecular Biology and Evolution, 16(1), 37–48. doi:10.1093/oxfordjournals.molbev.a026036

Ronquist, F., Teslenko, M., van der Mark, P., Ayres, D. L., Darling, A., Höhna, S., … Huelsenbeck, J. P. (2012). MrBayes 3.2: efficient Bayesian phylogenetic inference and model choice across a large model space. Systematic Biology, 61(3), 539–542. doi:10.1093/sysbio/sys029

Rousset, F. (2008). genepop’007: a complete re-implementation of the genepop software for Windows and Linux. Molecular Ecology Resources, 8(1), 103–106. doi:doi:10.1111/j.1471-8286.2007.01931.x

Schapira, A., & Boutsika, K. (2012). Malaria ecotypes and stratification. Advances in Parasitology, 78, 97–167. doi:https://doi.org/10.1016/B978-0-12-394303-3.00001-3

Schultner, E., Saramäki, J., & Helanterä, H. (2016). Genetic structure of native ant supercolonies varies in space and time. Molecular Ecology, 25(24), 6196–6213. doi:10.1111/mec.13912

Seto, K. C., Fragkias, M., Guneralp, B., & Reilly, M. K. (2011). A meta-analysis of global urban land expansion. PLoS One, 6(8), e23777. doi:10.1371/journal.pone.0023777

Shochat, E., Lerman, S. B., Anderies, J. M., Warren, P. S., Faeth, S. H., & Nilon, C. H. (2010). Invasion, competition, and biodiversity loss in urban ecosystems. BioScience, 60(3), 199–208. doi:10.1525/bio.2010.60.3.6

Shochat, E., Warren, P. S., Faeth, S. H., McIntyre, N. E., & Hope, D. (2006). From patterns to emerging processes in mechanistic urban ecology. Trends in Ecology & Evolution, 21(4), 186–191. doi:https://doi.org/10.1016/j.tree.2005.11.019

Souza, E., Follett, P. A., Price, D. K., & Stacy, E. A. (2008). Field suppression of the invasive ant Wasmannia auropunctata (Hymenoptera: Formicidae) in a tropical fruit orchard in Hawaii. Journal of Economic Entomology, 101(4), 1068–1074. doi:10.1093/jee/101.4.1068

Steiner, F. M., Schlick-Steiner, B. C., Moder, K., Stauffer, C., Arthofer, W., Buschinger, A., … Crozier, R. H. (2007). Abandoning aggression but maintaining self-nonself discrimination as a first stage in ant supercolony formation. Current Biology, 17(21), 1903–1907. doi:10.1016/j.cub.2007.09.061

Stow, A. J., Sunnucks, P., Briscoe, D. A., & Gardner, M. G. (2001). The impact of habitat fragmentation on dispersal of Cunningham’s skink (Egernia cunninghami): evidence from allelic and genotypic analyses of microsatellites. Molecular Ecology, 10(4), 867–878. doi:https://doi.org/10.1046/j.1365-294X.2001.01253.x

Suarez, A. V., Tsutsui, N. D., Holway, D. A., & Case, T. J. (1999). Behavioral and genetic differentiation between native and introduced populations of the Argentine ant. Biological Invasions, 1(1), 43–53. doi:10.1023/A:1010038413690

Tamura, K., Peterson, D., Peterson, N., Stecher, G., Nei, M., & Kumar, S. (2011). MEGA5: Molecular evolutionary genetics analysis using maximum likelihood, evolutionary distance, and maximum parsimony methods. Molecular Biology and Evolution, 28(10), 2731–2739. doi:10.1093/molbev/msr121

Theodorou, P., Baltz, L. M., Paxton, R. J., & Soro, A. (2021). Urbanization is associated with shifts in bumblebee body size, with cascading effects on pollination. Evolutionary Applications, 14(1), 53–68. doi:https://doi.org/10.1111/eva.13087

Tsutsui, N. D., & Suarez, A. V. (2003). The colony structure and population biology of invasive ants. Conservation Biology, 17(1), 48–58.

Tsutsui, N. D., Suarez, A. V., & Grosberg, R. K. (2003). Genetic diversity, asymmetrical aggression, and recognition in a widespread invasive species. Proceedings of the National Academy of Sciences, USA, 100(3), 1078. doi:10.1073/pnas.0234412100

Tsutsui, N. D., Suarez, A. V., Holway, D. A., & Case, T. J. (2000). Reduced genetic variation and the success of an invasive species. Proceedings of the National Academy of Sciences, USA, 97(11), 5948–5953. doi:10.1073/pnas.100110397

Vásquez, G. M., Schal, C., & Silverman, J. (2008). Cuticular hydrocarbons as queen adoption cues in the invasive Argentine ant. Journal of Experimental Biology, 211(8), 1249. doi:10.1242/jeb.017301

Vonshak, M., & Gordon, D. M. (2015). Intermediate disturbance promotes invasive ant abundance. Biological Conservation, 186, 359–367. doi:https://doi.org/10.1016/j.biocon.2015.03.024

Wang, J. (2011). COANCESTRY: a program for simulating, estimating and analysing relatedness and inbreeding coefficients. Molecular Ecology Resources, 11(1), 141–145. doi:doi:10.1111/j.1755-0998.2010.02885.x

Weir, B. S., & Cockerham, C. C. (1984). Estimating F-Statistics for the Analysis of Population Structure. Evolution, 38(6), 1358–1370. doi:10.2307/2408641

Wright, N. A., Steadman, D. W., & Witt, C. C. (2016). Predictable evolution toward flightlessness in volant island birds. Proceedings of the National Academy of Sciences, USA, 113(17), 4765. doi:10.1073/pnas.1522931113

Yakub, M., & Tiffin, P. (2017). Living in the city: urban environments shape the evolution of a native annual plant. Global Change Biology, 23(5), 2082–2089. doi:https://doi.org/10.1111/gcb.13528

Yang, C.-C., Ascunce, M. S., Luo, L.-Z., Shao, J.-G., Shih, C.-J., & Shoemaker, D. (2012). Propagule pressure and colony social organization are associated with the successful invasion and rapid range expansion of fire ants in China. Molecular Ecology, 21(4), 817–833. doi:doi:10.1111/j.1365-294X.2011.05393.x

Zheng, C., Yang, F., Zeng, L., Vargo, E. L., & Xu, Y. (2018). Genetic diversity and colony structure of Tapinoma melanocephalum on the islands and mainland of South China. Ecology and Evolution. doi:10.1002/ece3.4065

Zima, J., Lebrasseur, O., Borovanska, M., & Janda, M. (2016). Identification of microsatellite markers for a worldwide distributed, highly invasive ant species Tapinoma melanocephalum (Hymenoptera: Formicidae). European Journal of Entomology, 113, 409–414. doi:10.14411/eje.2016.053

