## Supplementary figures and tables for "Consistent signatures of urban adaptation in a native, urban invader ant *Tapinoma sessile*"

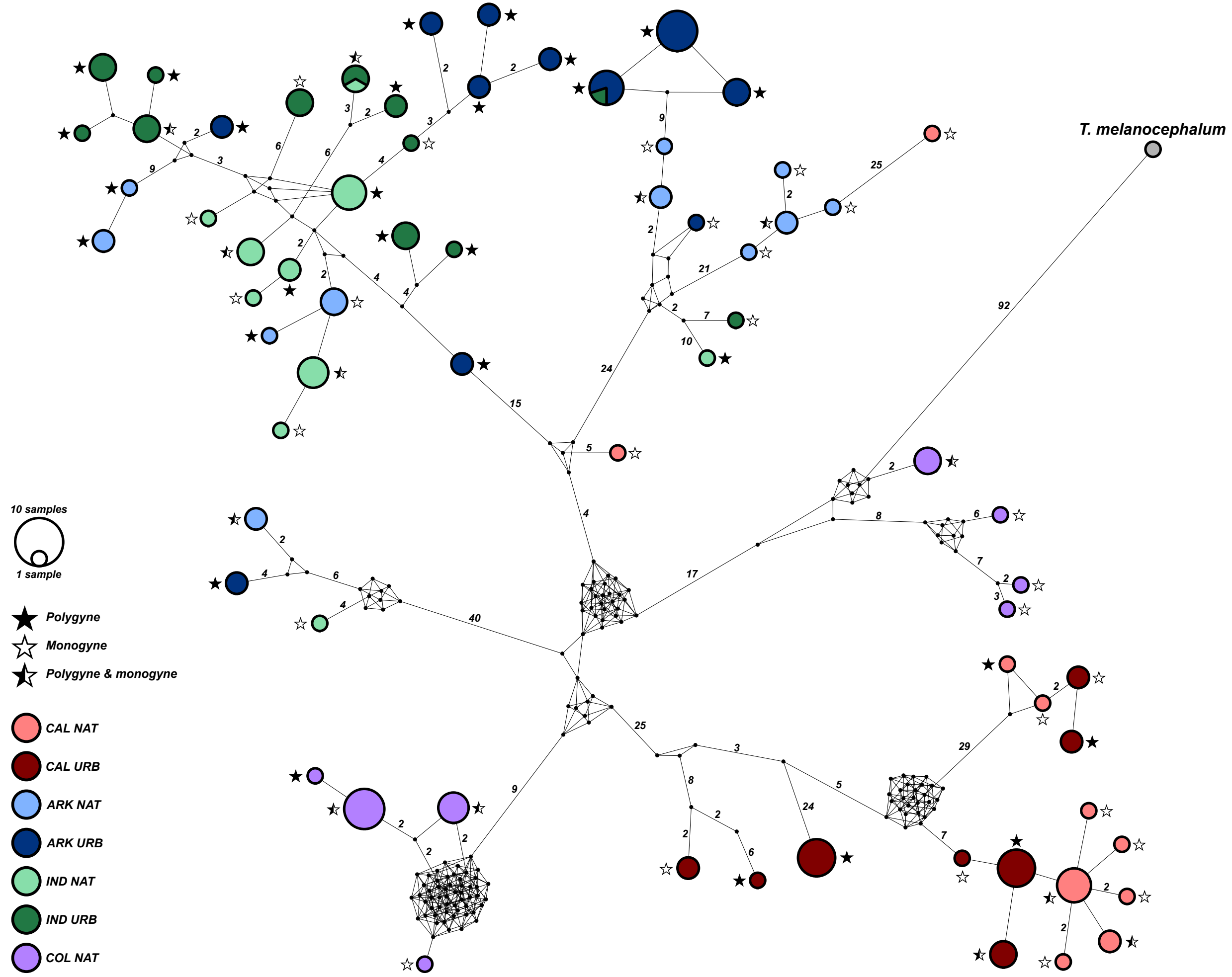


**Figure S1:** Median-joining haplotype network based on 145 COI sequences of *T. sessile* across the four localities, with one *T. melanocephalum* sequeunce as an outgroup. The numbers indicate the number of base pair (bp) differences between two nodes (*i.e.*, haplotypes); when numbers are not present on a branch the bp difference is one. Branch lengths are not proportional to the number of bp differences. The black circles represent median vectors (*i.e.*, hypothesized sequences).

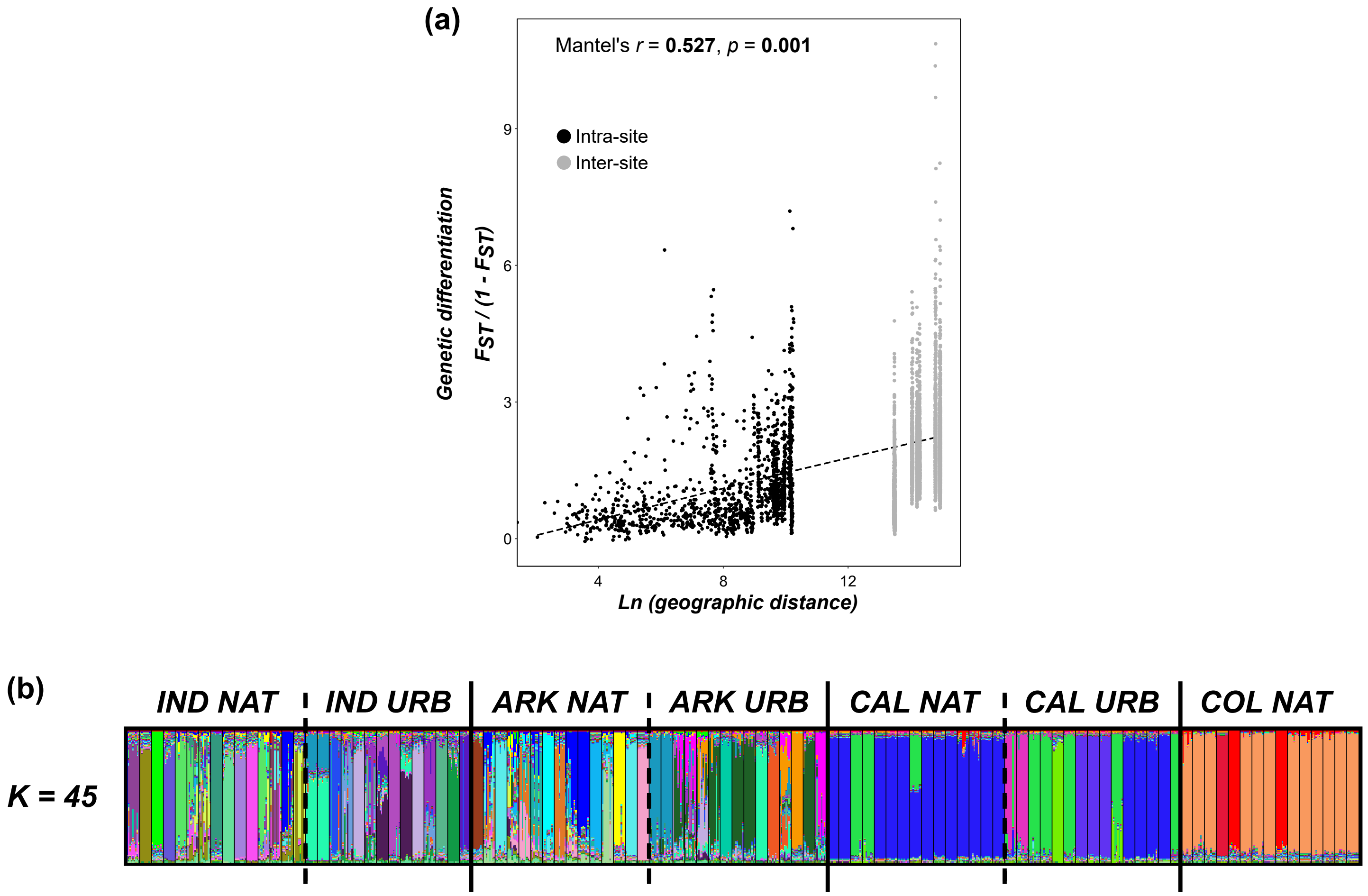


**Figure S2**: (a) Isolation-by-distance plot for each nest in the study. (b) STRUCTURE result for best K of the overall dataset (as determined by the Puechmaille method).

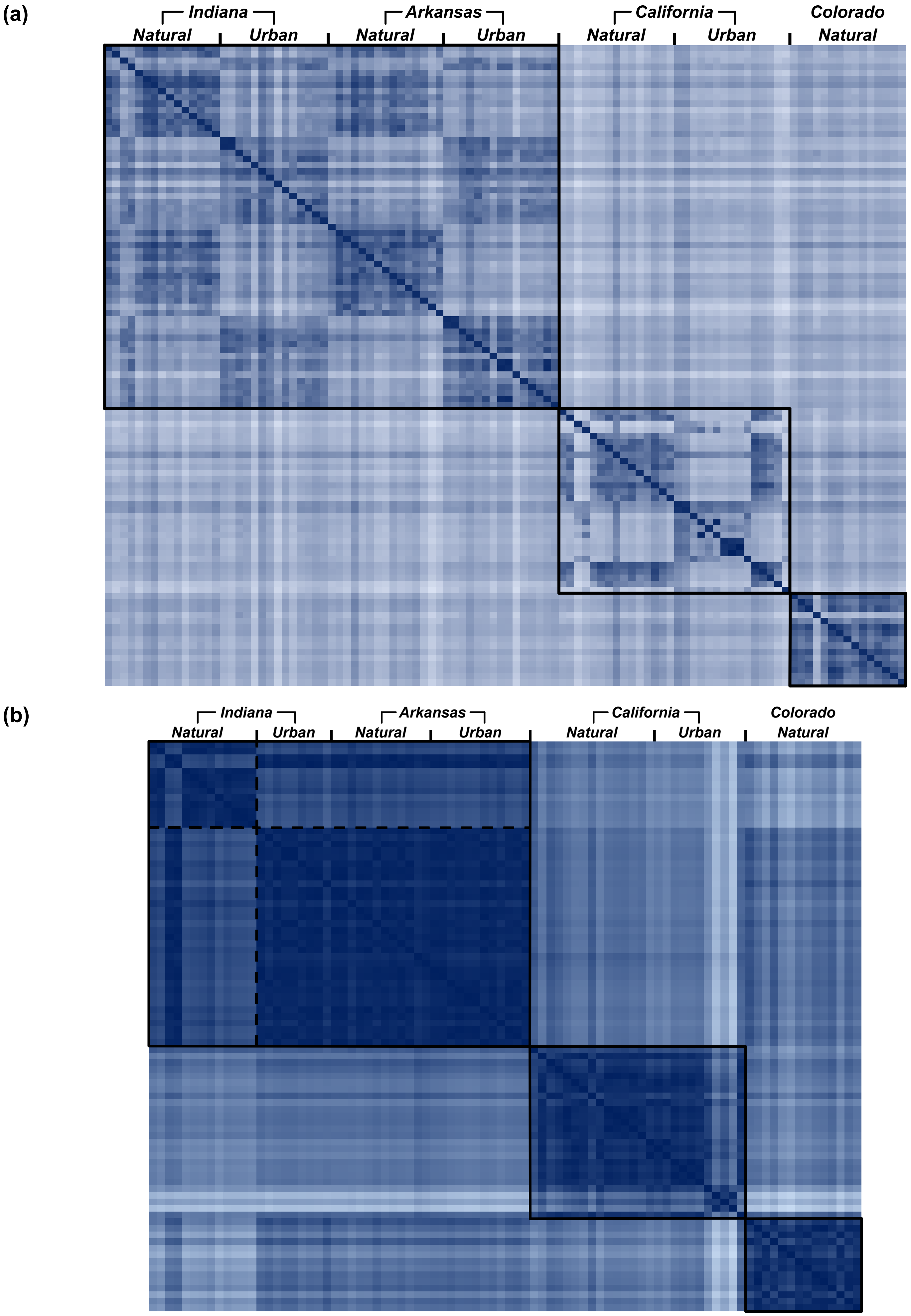


**Figure S3:** (a) Pairwise genetic differentiation ($F_{ST}$) between each nest sampled. Cells are shaded according to the genetic differentiation between a pair of nests, with darker and lighter regions indicative of lower and higher differentiation, respectively. (b) Pairwise chemical differentiation between each nest sampled. Cells are shaded according to the chemical differentiation between a pair of nests, with darker and lighter regions indicative of lower and higher differentiation, respectively.

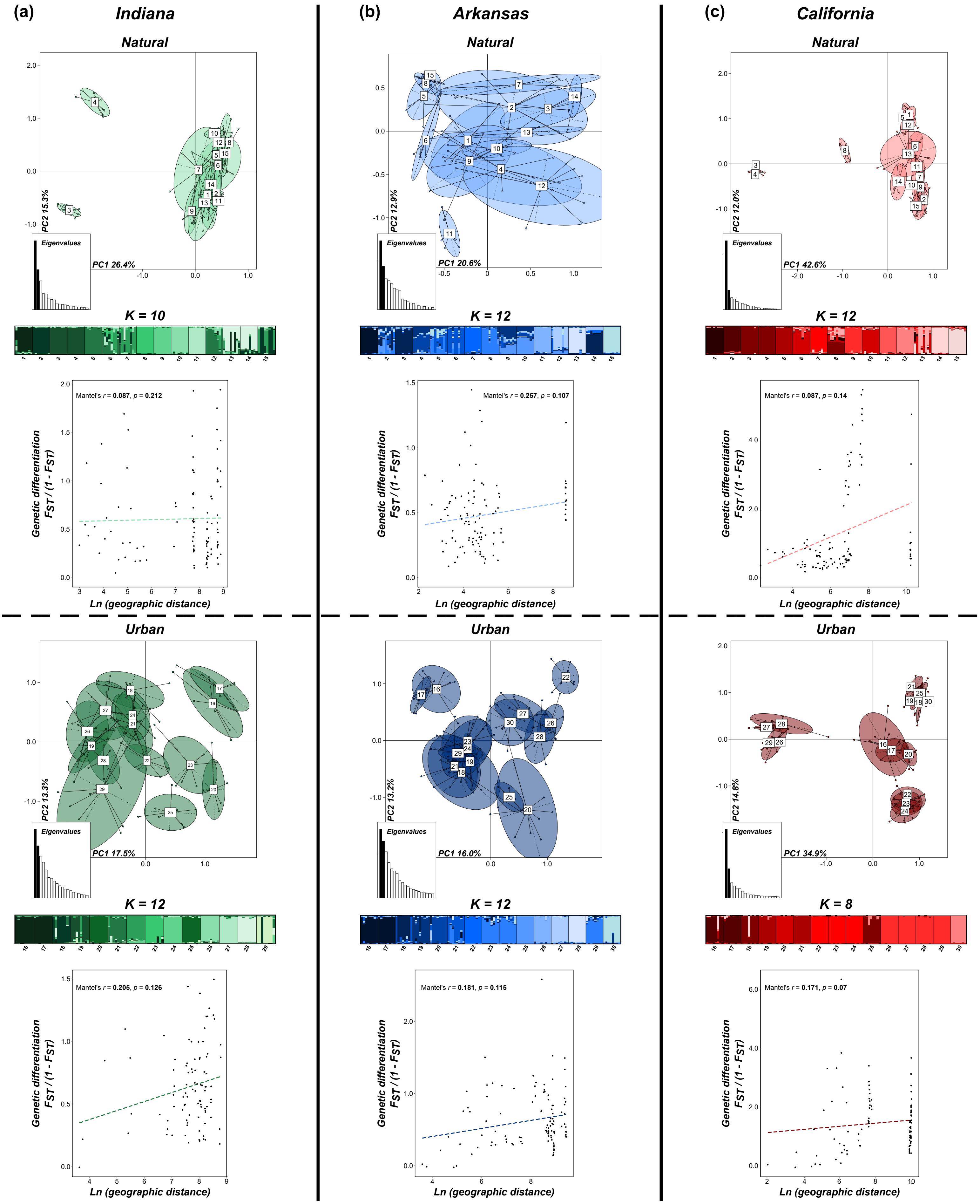


**Figure S4:** PCA, STRUCTURE and isolation-by-distance results for each habitat (*i.e.*, urban and natural) within (a) Indiana, (b) Arkansas and (c) California. By habitat results for Colorado are available in the main text as nests were only found in natural habitats there. For each STRUCTURE result, only the best *K* as determined by the Puechmaille method is shown.

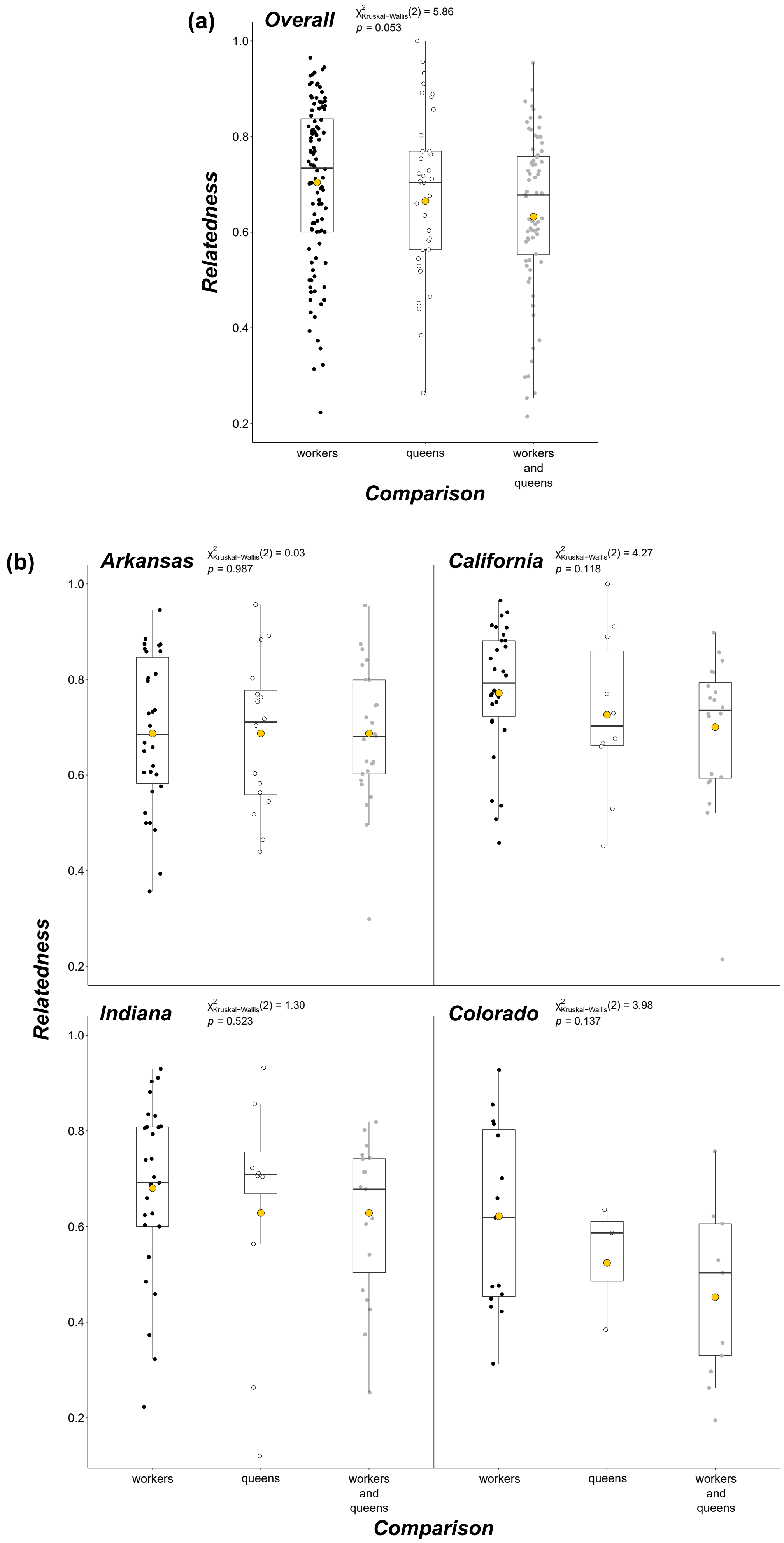


**Figure S5:** Comparisons of relatedness between workers, queens and both castes (a) overall and (b) within each locality. Each black, white and gray dot represents a single nest, and the gold dot on each boxplot denotes the mean of the group.

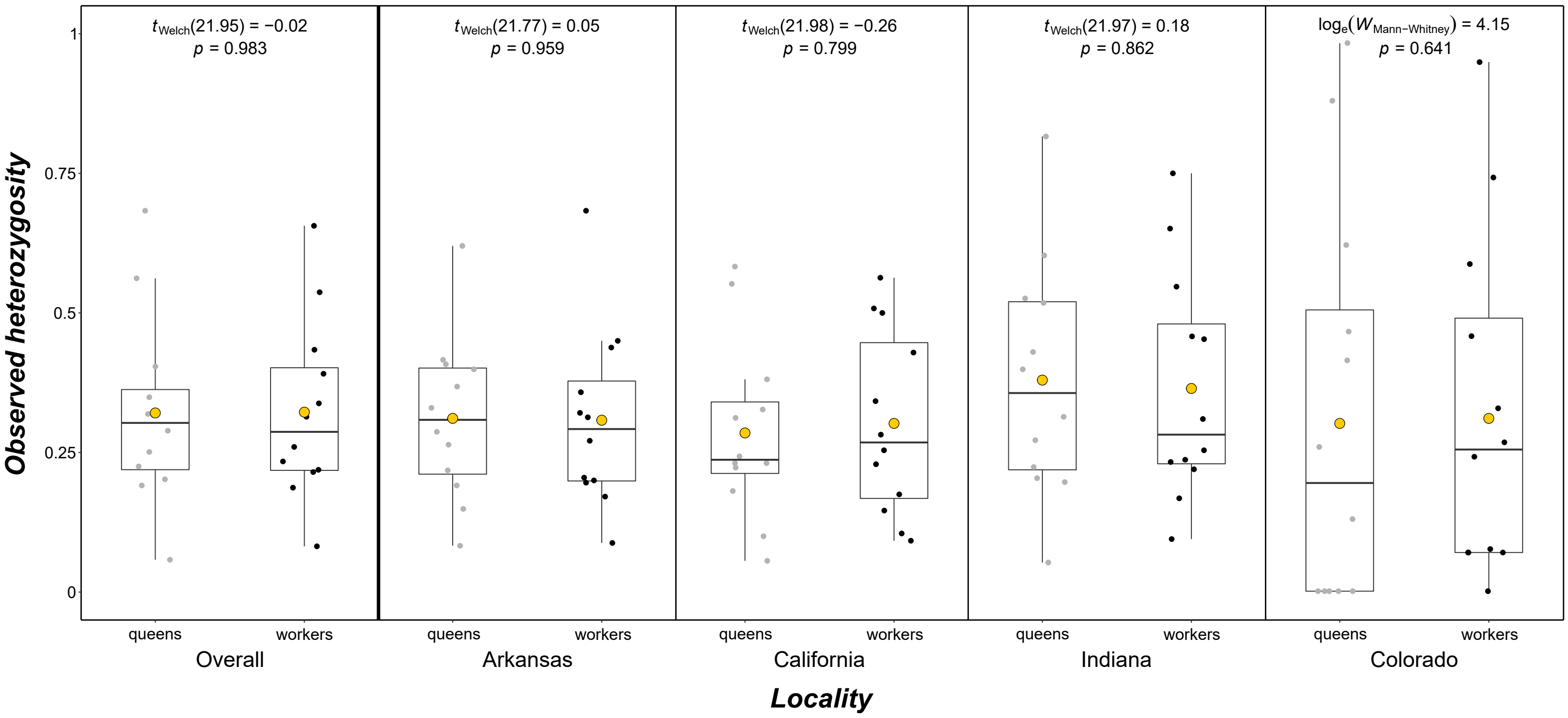


**Figure S6:** Comparisons of observed heterozygosity between queens and workers overall and across each of the four localities. Each black and gray dot represents the observed heterozygosity of a microsatellite marker (therefore, 12 dots per boxplot), and the gold dot on each boxplot denotes the mean of the caste.

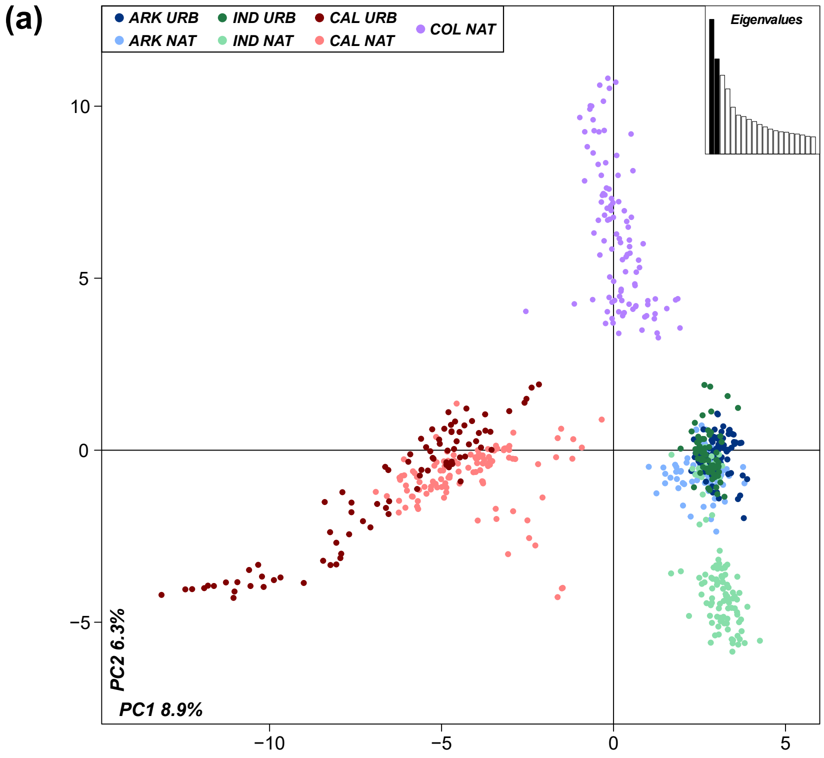


**Figure S7:** (a) PCA of the overall chemistry dataset. (b) Comparison of within nest chemical variation against the genetic diversity of the nest, in both the overall natural (left) and urban (right) dataset. The within nest chemical variation was calculated as the average Euclidean distance between each worker of a nest with the centroid of that nest (based on PC's 1 and 2 of the overall chemistry PCA). (c) Pairwise comparisons of chemical differentiation against geographic distance (left) and genetic differentiation (right) for the overall dataset. All significant comparisons are denoted with bolded *p*-values.

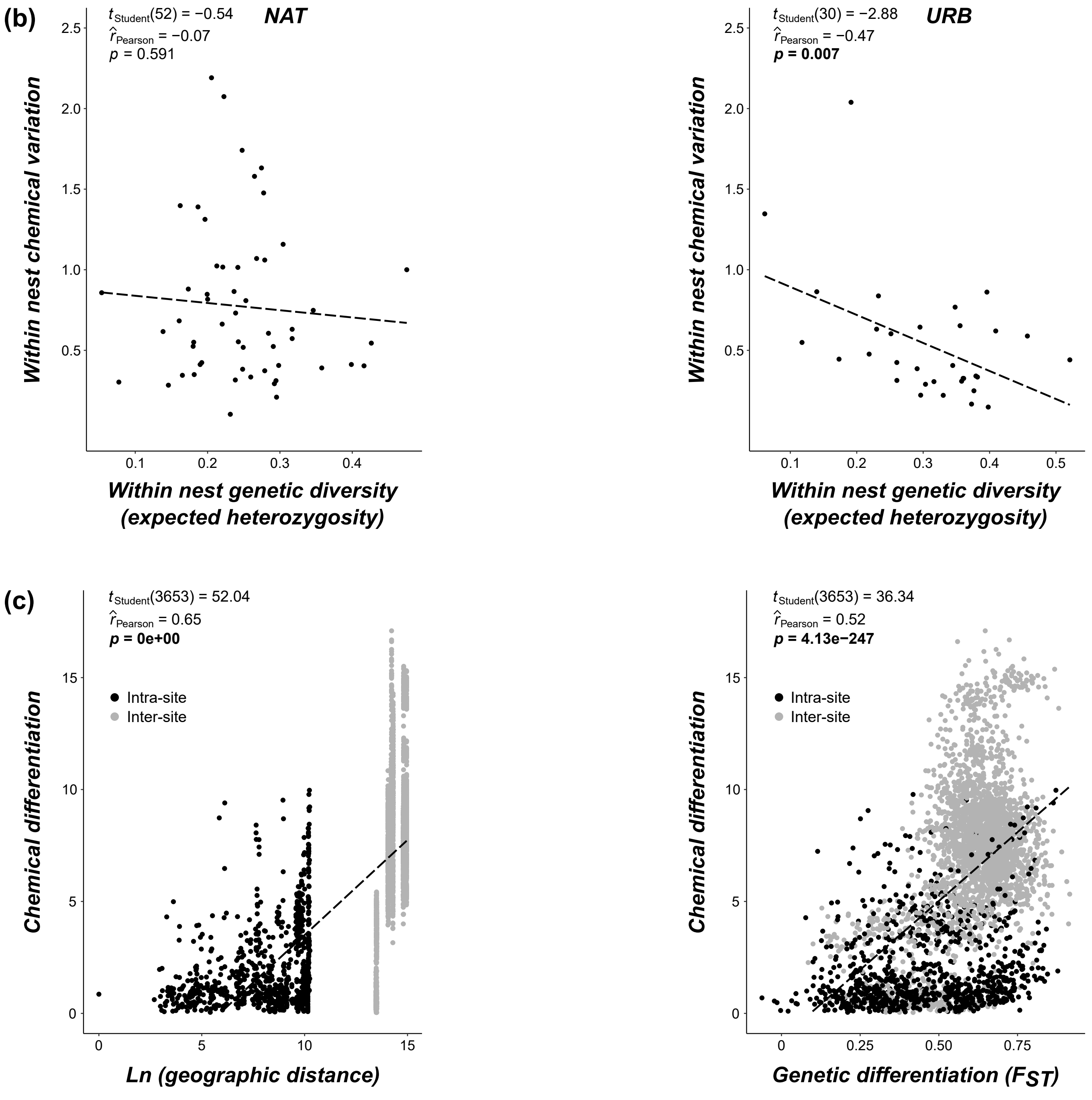

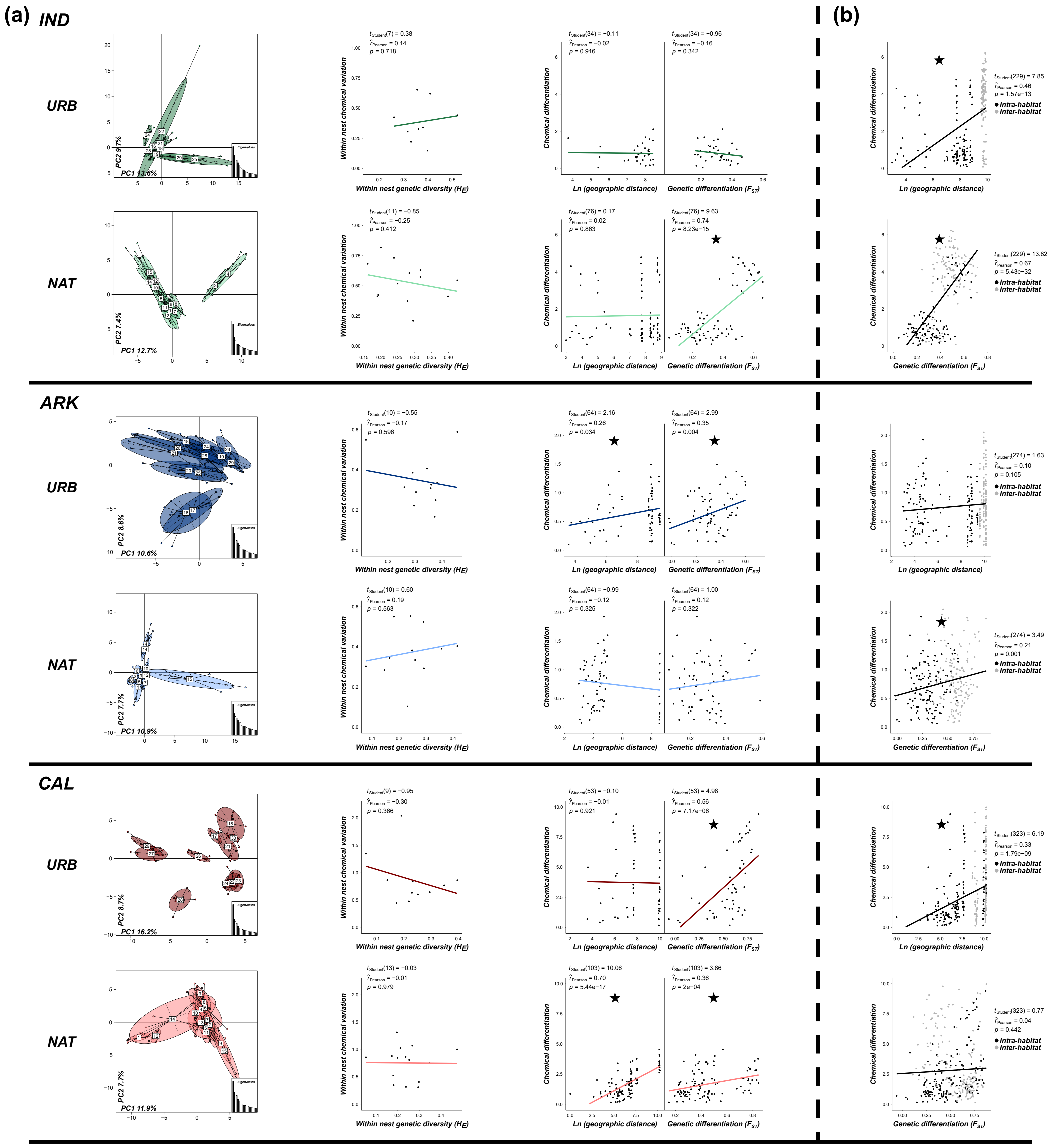


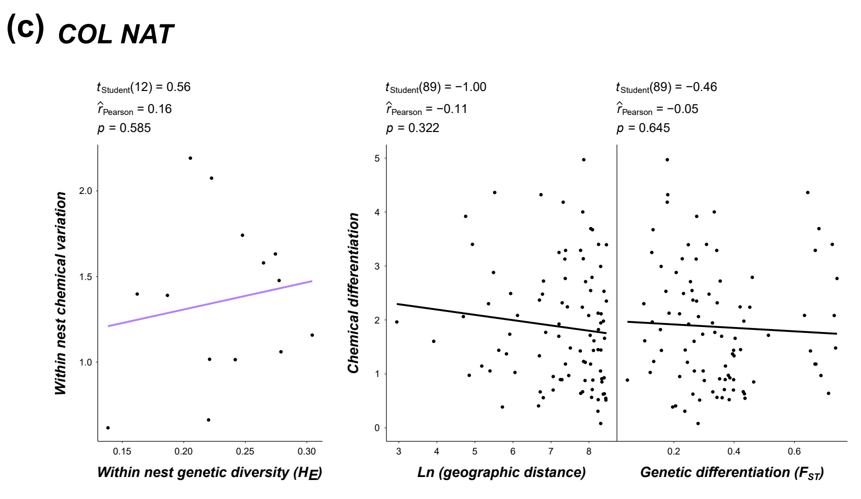


**Figure S8:** (a) Habitat-specific chemistry PCAs, correlations between chemical variation and genetic diversity, and correlations between chemical differentiation and 1) geographic distance and 2) genetic differentiation for each locality where nests were collected from multiple habitats. (b) Overall locality correlations (*i.e.*, both habitats combined) between chemical differentiation and 1) geographic distance and 2) genetic differentiation. (c) Correlations in the natural Colorado habitat. The PCA for this habitat can be seen in Figure 5d of the main text. Significant correlations are denoted with black stars.

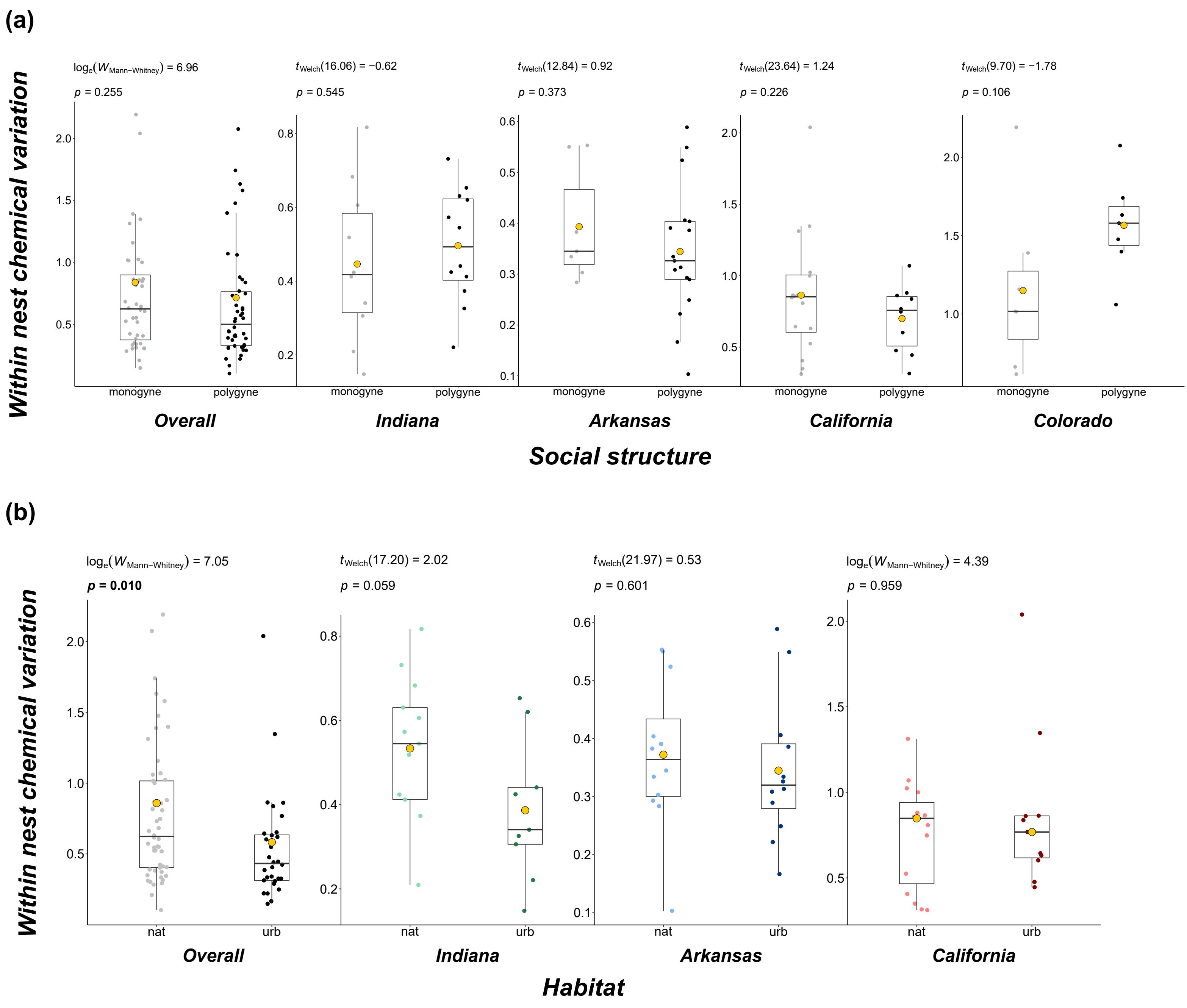


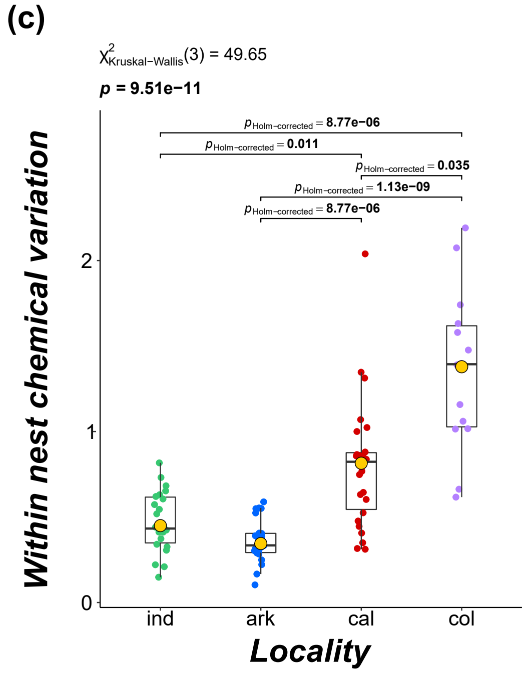


**Figure S9:** A comparison of within nest chemical variation means between (a) habitats, (b) social structures and (c) localities. The gold dot on each boxplot denotes the mean of the group, and bolded *p*-values indicate significance.

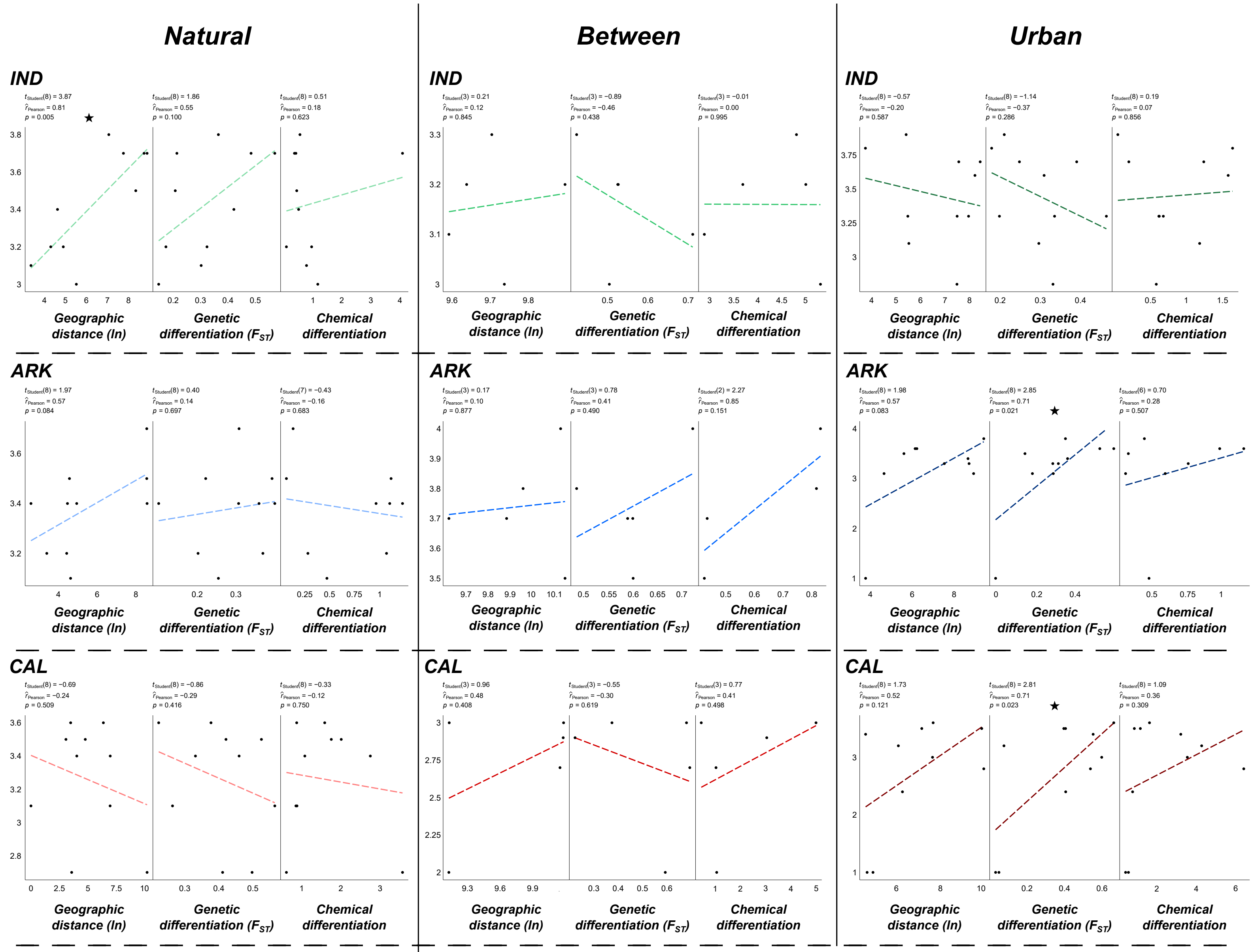


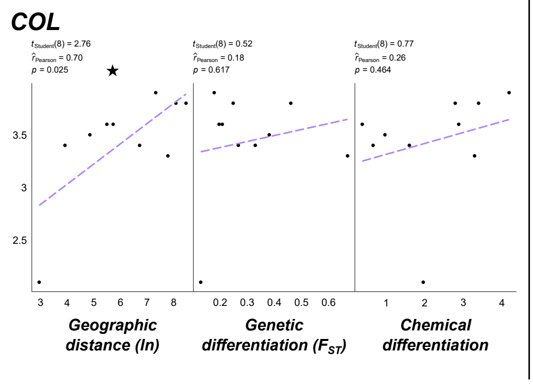


**Figure S10:** Correlations between aggression and 1) geographic distance, 2) genetic differentiation and 3) chemical differentiation. Each *y*-axis represents the mean aggression score between a pair of nests. Significant correlations are denoted with a black star on the plot.

**Table S1:** The collection location for each nest across the four localities. Nest numbers match the numbers in Figure 1.

| Locality | Habitat | Nest | Latitude | Longitude |
|---|---|---|---|---|
| Indiana | Natural | 1 | 39.180959 | -86.33819 |
| Indiana | Natural | 2 | 39.180869 | -86.338504 |
| Indiana | Natural | 3 | 39.179816 | -86.337378 |
| Indiana | Natural | 4 | 39.180516 | -86.33814 |
| Indiana | Natural | 5 | 39.180659 | -86.338393 |
| Indiana | Natural | 6 | 39.160625 | -86.346718 |
| Indiana | Natural | 7 | 39.160825 | -86.347721 |
| Indiana | Natural | 8 | 39.160838 | -86.34825 |
| Indiana | Natural | 9 | 39.161952 | -86.349601 |
| Indiana | Natural | 10 | 39.162509 | -86.349636 |
| Indiana | Natural | 11 | 39.118171 | -86.359556 |
| Indiana | Natural | 12 | 39.124975 | -86.349244 |
| Indiana | Natural | 13 | 39.123589 | -86.349895 |
| Indiana | Natural | 14 | 39.124507 | -86.349852 |
| Indiana | Natural | 15 | 39.12468 | -86.349793 |
| Indiana | Urban | 16 | 39.154215 | -86.536192 |
| Indiana | Urban | 17 | 39.153875 | -86.536199 |
| Indiana | Urban | 18 | 39.168817 | -86.537034 |
| Indiana | Urban | 19 | 39.171062 | -86.537287 |
| Indiana | Urban | 20 | 39.170549 | -86.536404 |
| Indiana | Urban | 21 | 39.155395 | -86.525399 |
| Indiana | Urban | 22 | 39.155838 | -86.522863 |
| Indiana | Urban | 23 | 39.139487 | -86.493143 |
| Indiana | Urban | 24 | 39.139119 | -86.517102 |
| Indiana | Urban | 25 | 39.141094 | -86.515968 |
| Indiana | Urban | 26 | 39.16706 | -86.494344 |
| Indiana | Urban | 27 | 39.164291 | -86.531262 |
| Indiana | Urban | 28 | 39.155467 | -86.566075 |
| Indiana | Urban | 29 | 39.155619 | -86.565622 |
| Arkansas | Natural | 1 | 34.844881 | -92.463909 |
| Arkansas | Natural | 2 | 34.840092 | -92.520507 |
| Arkansas | Natural | 3 | 34.841705 | -92.522486 |
| Arkansas | Natural | 4 | 34.839009 | -92.521296 |
| Arkansas | Natural | 5 | 34.839816 | -92.520902 |
| Arkansas | Natural | 6 | 34.839997 | -92.521072 |
| Arkansas | Natural | 7 | 34.840014 | -92.5214 |
| Arkansas | Natural | 8 | 34.839907 | -92.520454 |
| Arkansas | Natural | 9 | 34.839258 | -92.521248 |
| Arkansas | Natural | 10 | 34.839916 | -92.520157 |
| Arkansas | Natural | 11 | 34.840083 | -92.520362 |
| Arkansas | Natural | 12 | 34.840209 | -92.521116 |

| | | | | |
|---|---|---|---|---|
| Arkansas | Natural | 13 | 34.839952 | -92.521815 |
| Arkansas | Natural | 14 | 34.839951 | -92.521708 |
| Arkansas | Natural | 15 | 34.840009 | -92.520854 |
| Arkansas | Urban | 16 | 34.748422 | -92.399037 |
| Arkansas | Urban | 17 | 34.748096 | -92.399043 |
| Arkansas | Urban | 18 | 34.7225 | -92.34012 |
| Arkansas | Urban | 19 | 34.721875 | -92.340455 |
| Arkansas | Urban | 20 | 34.721506 | -92.341508 |
| Arkansas | Urban | 21 | 34.72493 | -92.339934 |
| Arkansas | Urban | 22 | 34.749524 | -92.320337 |
| Arkansas | Urban | 23 | 34.747946 | -92.26304 |
| Arkansas | Urban | 24 | 34.748356 | -92.26437 |
| Arkansas | Urban | 25 | 34.749052 | -92.269801 |
| Arkansas | Urban | 26 | 34.746894 | -92.265469 |
| Arkansas | Urban | 27 | 34.74651 | -92.28593 |
| Arkansas | Urban | 28 | 34.740941 | -92.271591 |
| Arkansas | Urban | 29 | 34.748742 | -92.264304 |
| Arkansas | Urban | 30 | 34.747232 | -92.27075 |
| California | Natural | 1 | 37.836961 | -122.487458 |
| California | Natural | 2 | 37.830895 | -122.184968 |
| California | Natural | 3 | 37.821498 | -122.163616 |
| California | Natural | 4 | 37.82745 | -122.172704 |
| California | Natural | 5 | 37.832505 | -122.183215 |
| California | Natural | 6 | 37.832271 | -122.182993 |
| California | Natural | 7 | 37.832271 | -122.182993 |
| California | Natural | 8 | 37.827344 | -122.172805 |
| California | Natural | 9 | 37.828458 | -122.175017 |
| California | Natural | 10 | 37.832076 | -122.181193 |
| California | Natural | 11 | 37.830683 | -122.185543 |
| California | Natural | 12 | 37.832025 | -122.180794 |
| California | Natural | 13 | 37.831723 | -122.180077 |
| California | Natural | 14 | 37.832695 | -122.182097 |
| California | Natural | 15 | 37.832601 | -122.181883 |
| California | Urban | 16 | 37.87323 | -122.274317 |
| California | Urban | 17 | 37.873299 | -122.274323 |
| California | Urban | 18 | 37.868624 | -122.261042 |
| California | Urban | 19 | 37.876987 | -122.269082 |
| California | Urban | 20 | 37.877177 | -122.272538 |
| California | Urban | 21 | 37.876556 | -122.269906 |
| California | Urban | 22 | 37.787008 | -122.466826 |
| California | Urban | 23 | 37.787485 | -122.468287 |
| California | Urban | 24 | 37.787271 | -122.467989 |
| California | Urban | 25 | 37.769113 | -122.470247 |
| California | Urban | 26 | 37.768853 | -122.472169 |

| | | | | |
|---|---|---|---|---|
| California | Urban | 27 | 37.766773 | -122.476428 |
| California | Urban | 28 | 37.76805 | -122.472988 |
| California | Urban | 29 | 37.768887 | -122.472592 |
| California | Urban | 30 | 37.769746 | -122.477634 |
| Colorado | Natural | 1 | 39.967879 | -105.266176 |
| Colorado | Natural | 2 | 39.967653 | -105.267426 |
| Colorado | Natural | 3 | 39.967482 | -105.267437 |
| Colorado | Natural | 4 | 39.986979 | -105.279117 |
| Colorado | Natural | 5 | 39.98785 | -105.280287 |
| Colorado | Natural | 6 | 39.987902 | -105.281785 |
| Colorado | Natural | 7 | 39.9880361 | -105.2842677 |
| Colorado | Natural | 8 | 39.995312 | -105.289634 |
| Colorado | Natural | 9 | 39.993972 | -105.295454 |
| Colorado | Natural | 10 | 39.993612 | -105.295809 |
| Colorado | Natural | 11 | 39.996263 | -105.296751 |
| Colorado | Natural | 12 | 39.997312 | -105.294556 |
| Colorado | Natural | 13 | 39.960376 | -105.267537 |
| Colorado | Natural | 14 | 39.959916 | -105.270309 |
| Colorado | Natural | 15 | 39.961031 | -105.2718 |

**Table S2:** The PCR primers and multiplexing arrangements.

| Name | M13 + Primer F | Primer R | PCR & Post-PCR Multiplex |
|---|---|---|---|
| Ant859 | CACGACGTTGTAAAACGACACTACGCGGAGAAACGTCTGGT | GTGATCTAAACTTCGATGAAC | F1 |
| Ant5035 | CACGACGTTGTAAAACGACACAGGATAGTTTCGCGGTTTATGG | ACTGACTCGYAGTGTATTTGAGGT | F1 |
| L911 | CACGACGTTGTAAAACGACACGCCTCGTCAAGAGTGGTCTC | GGAAAGCAGCAATTTTCTCG | N1 |
| Ant7249 | CACGACGTTGTAAAACGACACAAGTGTCAAGGGCGACTGAG | CGGGGACAATGGAGCAATCA | V1 |
| LP24 | CACGACGTTGTAAAACGACACGGAACAGGTGCTGAGAATCC | TGGCTAGTCCATGATTGTGC | V1 |
| Ant8424 | CACGACGTTGTAAAACGACACTCATAATGCAGATGATGGAACTCCT | GGCGAGTAACACAATGGCAC | P1 |
| Ant4155 | CACGACGTTGTAAAACGACACAGAATCTCTTGAGCCCGTCG | GGCGATACACTTCACCTGAGAC | F2 |
| Ant3653 | CACGACGTTGTAAAACGACACAGCAGAGACCAATCAACGGA | GGCAATTATCGGACCGGGTT | N2 |
| TM16 | CACGACGTTGTAAAACGACACAGCCAAGAGTTTCGTTCTCG | TCTCTTTACGACGGTTCGCT | N2 |
| Lhum19 | CACGACGTTGTAAAACGACACCTCTTAAAGCAATTGCATGTGG | ACGATCGCGTCCTTTGAG | V2 |
| Ant9218 | CACGACGTTGTAAAACGACACGACCCACTTTGCCCTCGTAA | CTCTCGATTAGTCAGGGTGGC | P2 |
| Ant3648 | CACGACGTTGTAAAACGACACCTCCTGGTCCTGGATCTCCA | TAACACCATGCCCTCTGTCG | P2 |

Ant* primer PCR conditions can be found in Butler, Siletti, Oxley, and Kronauer (2014); L911 primer conditions can be found in Zheng, Yang, Zeng, Vargo, and Xu (2018); LP24 primer conditions can be found in Berman, Austin, and Miller (2014); TM16 primer conditions can be found in Zima, Lebrasseur, Borovanska, and Janda (2016); lastly, Lhum19 primer conditions can be found in Krieger and Keller (1999). For the PCR & post-PCR multiplex, the letters correspond to the dye (6-FAM (F), NED (N), VIC (V) or PET (P)) and the numbers denote the post-PCR mix.

**References:**

Berman, M., Austin, C. M., & Miller, A. D. (2014). Characterisation of the complete mitochondrial genome and 13 microsatellite loci through next-generation sequencing for the New Caledonian spider-ant *Leptomyrmex pallens*. *Molecular Biology Reports, 41*(3), 1179-1187. doi:10.1007/s11033-013-2657-5

Butler, I. A., Siletti, K., Oxley, P. R., & Kronauer, D. J. (2014). Conserved microsatellites in ants enable population genetic and colony pedigree studies across a wide range of species. *PLOS One, 9*(9), e107334. doi:10.1371/journal.pone.0107334

Krieger, M. J. B., & Keller, L. (1999). Low polymorphism at 19 microsatellite loci in a French population of Argentine ants (*Linepithema humile*). *Molecular Ecology, 8*, 1078-1080.

Zheng, C., Yang, F., Zeng, L., Vargo, E. L., & Xu, Y. (2018). Genetic diversity and colony structure of *Tapinoma melanocephalum* on the islands and mainland of South China. *Ecology and Evolution*. doi:10.1002/ece3.4065

Zima, J., Lebrasseur, O., Borovanska, M., & Janda, M. (2016). Identification of microsatellite markers for a worldwide distributed, highly invasive ant species *Tapinoma melanocephalum* (Hymenoptera: Formicidae). *European Journal of Entomology, 113*, 409-414. doi:10.14411/eje.2016.053

**Table S3:** The number of base substitutions per locality from averaging over all sequence pairs within (bolded diagonal) and between (below diagonal) localities are shown. Analyses were conducted using the Kimura 2-parameter model involving 145 nucleotide sequences. All ambiguous positions were removed for each sequence pair (pairwise deletion option). There was a total of 658 positions in the final dataset.

| | Arkansas | California | Colorado | Indiana |
|---|---|---|---|---|
| **Arkansas** | **0.053 ± 0.006** | - | - | - |
| **California** | 0.094 ± 0.010 | **0.044 ± 0.005** | - | - |
| **Colorado** | 0.071 ± 0.008 | 0.087 ± 0.010 | **0.034 ± 0.005** | - |
| **Indiana** | 0.052 ± 0.006 | 0.098 ± 0.011 | 0.072 ± 0.009 | **0.025 ± 0.003** |

**Table S4:** The number and average number of alleles, observed ($H_O$) and expected ($H_E$) heterozygosity, inbreeding coefficient ($F_{IS}$) and fixation index ($F_{ST}$) for each of the 12 microsatellite loci.

| Marker | Alleles | Average | $H_O$ | $H_E$ | $F_{IS}$ | $F_{ST}$ |
|---|---|---|---|---|---|---|
| *Natural* | | | | | | |
| Ant859 | 8 | 1.67 | 0.263 | 0.218 | -0.208 | 0.705 |
| Ant5085 | 8 | 1.52 | 0.281 | 0.195 | -0.442 | 0.651 |
| Ant7249 | 19 | 2.78 | 0.719 | 0.517 | -0.389 | 0.362 |
| LP24 | 7 | 1.77 | 0.371 | 0.259 | -0.430 | 0.644 |
| L911 | 38 | 3.03 | 0.705 | 0.523 | -0.347 | 0.440 |
| Ant8424 | 4 | 1.42 | 0.206 | 0.150 | -0.374 | 0.663 |
| Ant4155 | 8 | 1.83 | 0.460 | 0.329 | -0.401 | 0.556 |
| Lhum19 | 3 | 1.22 | 0.124 | 0.086 | -0.449 | 0.726 |
| Ant3653 | 5 | 1.37 | 0.200 | 0.140 | -0.433 | 0.786 |
| Tm16 | 6 | 1.63 | 0.335 | 0.257 | -0.310 | 0.466 |
| Ant3648 | 6 | 1.67 | 0.352 | 0.231 | -0.525 | 0.605 |
| Ant9218 | 5 | 1.12 | 0.054 | 0.039 | -0.400 | 0.816 |
| *Urban* | | | | | | |
| Ant859 | 9 | 2.00 | 0.256 | 0.280 | 0.085 | 0.612 |
| Ant5085 | 3 | 1.25 | 0.135 | 0.098 | -0.377 | 0.802 |
| Ant7249 | 14 | 2.11 | 0.290 | 0.288 | -0.006 | 0.617 |
| LP24 | 9 | 2.34 | 0.418 | 0.430 | 0.029 | 0.474 |
| L911 | 37 | 3.84 | 0.589 | 0.627 | 0.055 | 0.345 |
| Ant8424 | 5 | 1.55 | 0.227 | 0.188 | -0.204 | 0.626 |
| Ant4155 | 6 | 2.16 | 0.399 | 0.369 | -0.083 | 0.462 |
| Lhum19 | 3 | 1.61 | 0.273 | 0.240 | -0.135 | 0.537 |
| Ant3653 | 5 | 1.79 | 0.281 | 0.304 | 0.070 | 0.571 |
| Tm16 | 6 | 1.91 | 0.341 | 0.361 | 0.057 | 0.375 |
| Ant3648 | 9 | 1.86 | 0.262 | 0.256 | -0.020 | 0.584 |
| Ant9218 | 4 | 1.32 | 0.120 | 0.093 | -0.277 | 0.844 |
| *Overall* | | | | | | |
| Ant859 | 11 | 1.81 | 0.260 | 0.244 | -0.066 | 0.677 |
| Ant5085 | 8 | 1.40 | 0.219 | 0.154 | -0.424 | 0.712 |
| Ant7249 | 20 | 2.50 | 0.537 | 0.420 | -0.279 | 0.500 |
| LP24 | 10 | 2.01 | 0.391 | 0.332 | -0.178 | 0.569 |
| L911 | 49 | 3.38 | 0.656 | 0.567 | -0.158 | 0.407 |
| Ant8424 | 5 | 1.47 | 0.215 | 0.166 | -0.293 | 0.645 |
| Ant4155 | 10 | 1.97 | 0.434 | 0.346 | -0.258 | 0.531 |
| Lhum19 | 3 | 1.38 | 0.187 | 0.151 | -0.237 | 0.651 |
| Ant3653 | 6 | 1.55 | 0.234 | 0.209 | -0.120 | 0.697 |
| Tm16 | 8 | 1.75 | 0.338 | 0.301 | -0.124 | 0.427 |
| Ant3648 | 9 | 1.75 | 0.314 | 0.242 | -0.300 | 0.669 |
| Ant9218 | 5 | 1.20 | 0.082 | 0.062 | -0.322 | 0.891 |

**Table S5:** Hierarchical partitioning of the genetic diversity assessed using an analysis of molecular variance (AMOVA) for the overall dataset.

| Source of variation | Sum of squares | Variance components | Percentage variation |
|---|---|---|---|
| Between localities | 2734.458 | 0.9226873 | 10.623174 |
| Between habitats within localities | 1600.92 | 2.0817376 | 23.967663 |
| Between nests within habitats | 3924.92 | 2.3644543 | 27.222666 |
| Between individuals within nests | 1946.697 | -0.6390175 | -7.357198 |
| Within individuals | 3287.226 | 3.9557474 | 45.543696 |
| Total | **13494.222** | **8.6856091** | |

**Table S6:** Hierarchical partitioning of the genetic diversity assessed using an analysis of molecular variance (AMOVA) in Arkansas, California and Indiana.

***Arkansas***

| Source of variation | Sum of squares | Variance components | Percentage variation |
|---|---|---|---|
| Between habitats | 811.2135 | 3.2527749 | 38.982381 |
| Between nests within habitats | 855.3301 | 1.7205692 | 20.619898 |
| Between individuals within nests | 633.8634 | -0.3524766 | -4.224202 |
| Within individuals | 893.6041 | 3.7233505 | 44.621922 |
| Total | **3194.0111** | **8.344218** | |

***California***

| Source of variation | Sum of squares | Variance components | Percentage variation |
|---|---|---|---|
| Between habitats | 313.3899 | 1.0666696 | 13.874517 |
| Between nests within habitats | 1636.0865 | 3.517233 | 45.749789 |
| Between individuals within nests | 500.0088 | -0.7116869 | -9.257143 |
| Within individuals | 911.9668 | 3.8157607 | 49.632837 |
| Total | **3361.4521** | **7.6879763** | |

***Indiana***

| Source of variation | Sum of squares | Variance components | Percentage variation |
|---|---|---|---|
| Between habitats | 475.3099 | 1.8912037 | 24.7244 |
| Between nests within habitats | 1000.9533 | 2.1387454 | 27.9606 |
| Between individuals within nests | 579.0405 | -0.7667736 | -10.02431 |
| Within individuals | 1017.5435 | 4.3859632 | 57.33931 |
| Total | **3072.8471** | **7.6491388** | |
